# The ferredoxin/flavodoxin-NADP^+^ oxidoreductase YumC is essential for isoprenoid and peptidoglycan biosynthesis in *Bacillus subtilis*

**DOI:** 10.64898/2026.09.04.749439

**Authors:** Deniz Akbulut, Marirene Chacon-Arnaude, Dillon P. McBee, Anne Lamsa, Alan I. Derman, Joshua A. Baccile, Javier Lopez-Garrido

## Abstract

Redox reactions mediated by ferredoxin/flavodoxin-NADP^+^ oxidoreductases (FNRs) and their associated electron-carrier proteins, ferredoxins and flavodoxins, are essential in biology. Although the biochemical activities of these redox proteins are conserved, their precise physiological roles can differ among organisms and cannot be easily inferred. Here we have defined an essential role for *Bacillus subtilis* YumC, a member of a distinct group of bacterial FNRs that resemble thioredoxin reductase. We have used targeted protein degradation, cytological profiling, metabolomics, and genetic complementation to show that YumC catalyzes the transfer of electrons from NADPH, through ferredoxin (Fer) or through the flavodoxin YkuP, to the isoprenoid biosynthesis pathway, and specifically to the redox enzyme IspG. When YumC was degraded, isoprenoid biosynthesis was compromised, and the level of undecaprenyl phosphate, the isoprenoid lipid carrier for peptidoglycan building block translocation, was diminished. Degradation of YumC or of Fer in a Δ*ykuP* strain led to defective peptidoglycan biosynthesis, activation of the σ^M^-dependent cell-wall stress response, and lethality. The introduction into *B. subtilis* of an alternative pathway for isoprenoid biosynthesis that does not require input from electron-carrier proteins could complement the degradation of Fer in a Δ*ykuP* strain, but not the degradation of YumC. This finding indicates that YumC is required for other essential processes that do not necessarily involve Fer and YkuP. This work provides an explanation for why YumC is essential, reveals how reducing power is delivered to isoprenoid biosynthesis in *B. subtilis,* and illustrates how the varied roles of redox systems among bacteria depend upon metabolic context.

## INTRODUCTION

The movement of electrons is an integral part of metabolism (1). Redox reactions, in which electrons are transferred from one chemical species to another, account for roughly one-third of known enzymatic reactions (2). Electrons from oxidation reactions can be captured as NADH and NADPH, which provide reducing equivalents for enzymatic reactions and for the generation of energy. Many enzymes can be reduced directly by NADH or NADPH, but others rely on intermediary electron-carrier proteins (ECPs) such as ferredoxins and flavodoxins to shuttle electrons between NAD(P)H and redox enzymes. The transfer of electrons via ECPs is mediated by prosthetic groups within the protein, by iron-sulfur clusters in the ferredoxins, and by a flavin mononucleotide (FMN) in the flavodoxins. Ferredoxin and flavodoxins are ubiquitous in biology (3), and they participate in a wide variety of key metabolic processes, including photosynthesis, nitrogen fixation, biotin biosynthesis, and lipid metabolism (4, 5).

Ferredoxin (flavodoxin)-NADP^+^ oxidoreductases (FNRs) are the enzymes that catalyze the transfer of electrons from NADPH to the ECPs (6). FNRs contain flavin adenine dinucleotide (FAD) as a prosthetic group, which is reduced to the hydroquinone state when it accepts an electron pair from NADPH. The FAD can donate these electrons one by one to the one-electron acceptor ECPs, as it transits through the radical semiquinone and oxidized hydroquinone states (7). This chemistry allows the two-electron hydride of NADPH to support reactions that proceed through single-electron transfers. During the light reactions of photosynthesis, the system runs in reverse, and electrons from Photosystem I reduce ferredoxin, and the FNR catalyzes the reduction of NADP^+^ to NADPH with electrons from ferredoxin. The FNRs comprise several phylogenetic groups. The most recently characterized of these consists of homodimeric proteins that closely resemble thioredoxin reductase but lack their critical cysteine-containing motifs (8, 9). Within this group are the FNRs of the Gram-positive Firmicutes such as *Bacillus subtilis* (9). *B. subtilis* has two FNR genes, *yumC*, which is essential, and *ycgT*, which is not (10). YumC was first purified twenty years ago (9), and its crystal structure solved a few years later (11). YumC has been the subject of extensive biochemical and biophysical study (9, 12–15). YcgT has received less attention, although it appears to be able to function as an FNR (16).

Because *yumC* is essential, YumC would be presumed to catalyze the transfer of electrons from NADPH through the *B. subtilis* ferredoxin Fer, or through its flavodoxins YkuN and YkuP, to at least one enzyme that is essential for the viability of the cell. The *ykuN* and *ykuP* genes are cotranscribed in an operon that also contains *ykuO*, a nonessential gene of unknown function. The *yumC* and *fer* genes are each monocistronic and they are not linked to each other or to *ykuNOP*. *yumC* is essential, but *fer*, *ykuN*, and *ykuP* are each individually dispensable (10, 17). Preventing transcription of the *ykuNOP* operon in a strain containing a disruption of the *fer* gene leads to the cessation of growth, indicating that either YkuN, YkuO, YkuP or some combination of these three proteins is required when Fer is not present. There is, however, no phenotype when transcription of *ykuNOP* is prevented in a Fer^+^ strain, indicating that all three proteins are dispensable so long as Fer is present (18). The simple conclusion to be drawn from these observations is that the essential function of YumC requires at least one ECP, either Fer, or one or possibly both of the flavodoxins. But to date, no essential enzyme or metabolic process has been identified in *B. subtilis* that requires either YumC or both YumC and an ECP.

Several studies have implicated YumC or an ECP in a specific cellular reaction or pathway. Both *fer* and *ykuNOP* were required *in vivo* for a wild-type level of lipid desaturase activity from the enzyme Δ5-Des in *B. subtilis.* The FNR was not identified (18). Biochemical studies have typically paired YumC with ECPs from other organisms, or FNRs from other organisms with *B. subtilis* ECPs. Both YkuN and YkuP could reduce the heme in *B. subtilis* nitric oxide synthase (bNOS), YkuN being the more efficient, and an FNR was required; this study used an FNR from *E. coli* (19). When YumC was used, YkuN could support bNOS activity efficiently only when it was fused to bNOS (20), and a triparte YumC-YkuN-bNOS fusion also showed enhanced activity (21). Fer, YkuN, or YkuP supported lipid hydroxylation by the enzyme BioI during biotin biosynthesis when paired with an FNR from *E. coli* (22, 23). The enzymatic activity of the cytochrome P450 109B1 monooxygenase (CYP109B1; YjiB) and that of the radical SAM 7-carboxy-7-deazaguanine synthase (QueE) could be reconstituted with YkuN or YkuP along with an *E. coli* FNR (24, 25). Although none of these enzymes are essential in *B. subtilis* (10), these studies suggest that YumC and the *B. subtilis* ECPs could form a functional redox module *in vivo*, with Fer, YkuN, and YkuP relaying electrons to enzymes in a variety of different processes (Fig. 1A).

**Fig. 1.**
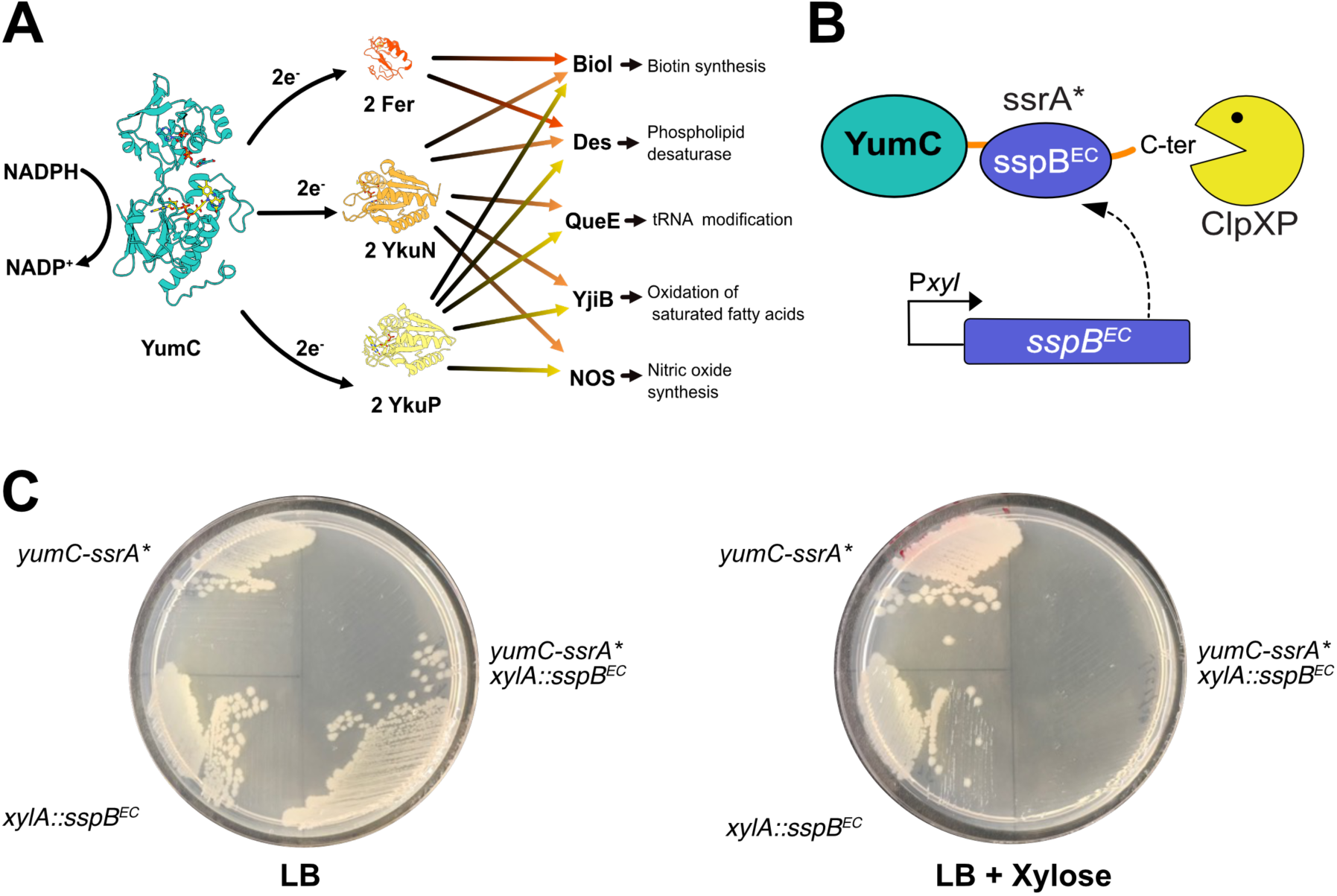
YumC is essential for the viability of *B. subtilis*. (A) YumC, a ferredoxin (flavodoxin)-NADP^+^ oxidoreductase (FNR), catalyzes the transfer of electrons from NADPH via a non-covalently bound FAD to ferredoxin (Fer) or to a flavodoxin (YkuN or YkuP), and these electron carrier proteins (ECPs) provide electrons to various cellular reactions. Ribbon diagram of YumC adapted from PDB 3LZW. BioI, cytochrome P450 CYP107H1; Des, phospholipid desaturase; QueE, 7-carboxy-7-deazaguanine synthase; YjiB, monooxygenase cytochrome P450 CYP109B1. Pairings of ECPs with oxidizing substrates is based on biochemical or physiological studies. See main text for details. (B) Regulated intracellular degradation of a target protein. The protein (here YumC) tagged with ssrA* at its C-terminus is degraded when conveyed to the endogenous ClpXP protease by SspB^EC^. Proteolysis is triggered by the induction of *sspB*^EC^ with xylose. (C) Assessment of the requirement for YumC. Strains were streaked onto LB agar plates containing no xylose or 1% xylose, and incubated for 18 hours at 30 °C (upper left, JLG1680; lower left, JLG364; right, JLG1728). The same outcome was obtained at 37 °C.

The essentiality of YumC in *B. subtilis* has made its physiological function difficult to dissect *in vivo*. The TrxB2 protein of *Lactococcus lactis*, which is a YumC-like FNR, is required for aerobic growth and this requirement was linked to the activity of the class Ib ribonucleotide reductase (26). *L. lactis trxB2* mutants barely grow aerobically, but this defect could be eliminated by providing exogenous deoxyribonucleosides and could be suppressed by specific mutations in *nrdI* (26). The *nrdI* gene codes for a flavodoxin that is required for the generation of the tyrosyl radical in the β subunit (NrdF) of class Ib ribonucleotide reductases (27). YumC-like FNRs have also been shown to reduce *B. cereus* NrdI *in vitro* (28). These findings raise the possibility that YumC provides electrons directly to the *B. subtilis* NrdI and is therefore required for ribonucleotide reduction. *B. subtilis* has only one ribonucleotide reductase, it is class Ib, and *nrdI* is essential (17). But YumC is required in *B. subtilis* even when its growth medium is supplemented with deoxyribonucleosides (26).

Here, we demonstrate that YumC is essential in *B. subtilis* for the biosynthesis of isoprenoids via the endogenous methylerythritol (MEP) pathway. YumC functions with Fer or the flavodoxin YkuP to provide electrons to the redox reaction catalyzed by the isoprenoid biosynthetic enzyme IspG and likely also to the redox reaction catalyzed by IspH. In the absence of YumC, the cellular pool of the isoprenoid undecaprenyl phosphate (Und-P) is depleted. Und-P is the lipid carrier that mediates the translocation of the cytoplasmic peptidoglycan (PG) building block across the cytoplasmic membrane for its incorporation into the cell wall. Its depletion leads to disruption of PG synthesis and cell lysis. Introduction of the alternative mevalonate (MVA) isoprenoid biosynthetic pathway, which does not require electrons from ECPs, complements the degradation of Fer in cells lacking YkuP, but does not complement the degradation of YumC. YumC must therefore also be essential for other yet to be discovered process(es).

## RESULTS

### YumC is essential in *Bacillus subtilis* for both aerobic and anaerobic growth

We first confirmed that YumC is essential by degrading the protein in the cell. We tagged *yumC* at its native chromosomal locus with a modified *E. coli* ssrA (ssrA*) degradation tag, and we triggered degradation of YumC by producing the cognate *E. coli* adaptor SspB (SspB^EC^), which recognizes the tag and delivers the protein to the endogenous *B. subtilis* protease ClpXP for degradation (Fig. 1B) (29). We induced degradation of YumC-ssrA* by expressing *sspB^EC^*from a xylose-inducible promoter. *B. subtilis* strains carrying both *yumC-ssrA\** and the xylose-inducible *sspB^EC^* failed to grow in the presence of xylose, whereas control strains containing only *yumC-ssrA\** or only *sspB^EC^* grew normally (Fig. 1C). The *ycgT* gene codes for a YumC paralog that is 47% identical to YumC at the amino acid level, but is not essential (10). *ycgT* is part of the *fur* regulon and its transcription is derepressed roughly tenfold in response to iron starvation (30). Expression of the *B. subtilis* 168 *ycgT* gene from an IPTG-inducible hyper-spank promoter failed to restore viability when YumC-ssrA* was degraded (Fig. S1).

The *L. lactis* YumC-like reductase TrxB2 is required for aerobic growth but not for anaerobic growth (26), so we determined whether YumC was dispensable when *B. subtilis* was grown anaerobically (31, 32). *B. subtilis* grew robustly under anaerobic conditions in LB plates supplemented with arabinose, with nitrate serving as terminal electron acceptor, but failed to grow at all when YumC was degraded (Fig. S2). YumC is therefore essential in *B. subtilis* in both aerobic and anaerobic environments, so its functions likely differ from or exceed the functions of *L. lactis* TrxB2, as has been proposed (26).

### Suppressors of YumC depletion could not be isolated

We employed two genetic selections to reveal the function of YumC (see Supplementary Methods for details). From the first selection, in which we simply characterized several of the many spontaneous mutants that arose when YumC was degraded, we recovered only mutations that would be expected to interfere with the degradation of YumC, for example mutations in *clpX* or mutations that altered the ssrA* tag. For the second selection, we again degraded YumC, but we also restricted its production. We deleted the native *yumC* gene from the chromosome and we constructed an operon under control of the IPTG-inducible hyper-spank promoter that consisted of *lacZ* followed by *yumC*-*ssrA*\*; this strain also contained *lacI*, and was therefore viable only in the presence of IPTG. *sspB^EC^* was present in the strain, under the control of the xylose-inducible promoter (Fig. S3A, upper diagram). The selection plates lacked IPTG, so as to prevent expression of *yumC-ssrA\**, and they also contained xylose, so as to induce SspB^EC^-dependent degradation of any YumC-ssrA* that did happen to be produced (Fig. S3A, lower diagram). Because the plates contained X-Gal as well, we were able to exclude from further investigation any blue colonies from among those that arose spontaneously, as these would have been derived from mutations that led to the derepression of the synthetic operon. But we recovered among the white colonies only mutations that amplified *yumC*-*ssrA\**, up to 30 times, which was presumably sufficient to outpace the cell’s allotment of both Lac repressor and SspB^EC^, and thereby allow for the maintenance of an adequate steady-state level of YumC (Fig. S3B). The inability to recover genuine suppressor mutations that could compensate for the loss of or circumvent the requirement for YumC underscored that YumC is indeed essential in *B. subtilis* and also suggested that the protein may be required at more than one stage in an essential process or in more than one essential process.

### YumC is required for cell wall biosynthesis

We were able to identify one essential process for which YumC is required with Rapid Inhibition Profiling (RIP) (33, 34). RIP couples the rapid inducible degradation of ssrA*-tagged proteins with Bacterial Cytological Profiling (35) to monitor cytological changes that arise in connection with target protein degradation (Fig. 2A). Degradation of proteins that are required for an essential cellular process, for example transcription, DNA replication, or PG biosynthesis, gives rise to a distinct set of cytological changes that taken together constitute the bacterial cytological profile (Fig. 2B). The same profile is generated by antibiotics that target the same cellular process (33). The library of cellular processes and the associated cytological profiles that their disruptions produce can then be used to identify the process in which an uncharacterized essential protein is involved.

**Fig. 2.**
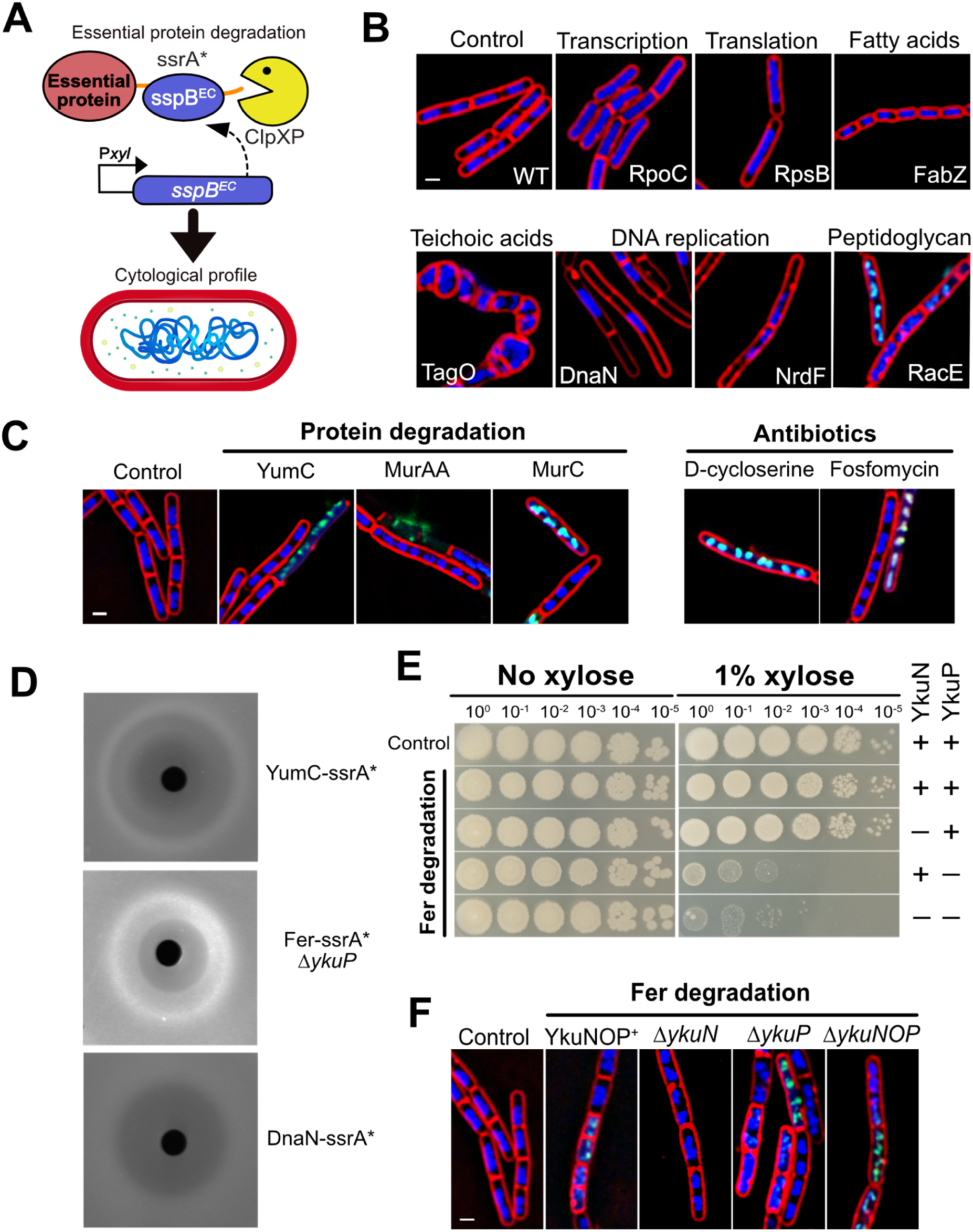
Degradation of YumC and associated ECPs leads to peptidoglycan biosynthesis defects. (A) Rapid Inhibition Profiling (RIP) entails the rapid, inducible degradation of essential proteins, followed by the assessment of the resulting cytological profile by fluorescence microscopy. (B) RIP of essential proteins in different pathways produce distinct profiles. The ssrA*-tagged protein subjected to degradation is indicated within each micrograph, and the pathway affected on top. Membranes were stained with FM4-64 (red), and DNA was stained with DAPI (blue) and with SYTOX Green (green), which is ordinarily cell-impermeant but enters the cell when envelope permeability is compromised. Scale bar, 1 µm. All strains contain *xylA*::*sspB^EC^*: WT, JLG364; *rpoC-ssrA\**, JLG7235; *rpsB-ssrA\**, JLG7234; *fabZ-ssrA\**, JLG707; *tagO-ssrA\**, JLG1170; *dnaN-ssrA\**, JLG715; *nrdF-ssrA\**, JLG1251; *racE-ssrA\**, JLG1245. (C) RIP of YumC and of essential proteins in the peptidoglycan biosynthetic pathway. Cultures were grown in LB medium, treated with 1% xylose to trigger degradation of the ssrA*-tagged protein (indicated on top of each micrograph), or with antibiotics that inhibit peptidoglycan biosynthesis, and imaged on agarose pads during exponential phase. Scale bar, 1 μm; all panels are at the same scale. All strains contain *xylA*::*sspB^EC^*: control, JLG364; *yumC*-*ssrA\**, JLG1728; *murAA*-*ssrA\**, JLG736; *murC*-*ssrA\**, JLG380. D-cycloserine and fosfomycin were added to a final concentration of 47.9 μg/mL and 187.5 μg/mL respectively. (D) Induction of the σ^M^ cell wall stress response upon degradation of different target proteins, assessed by disk diffusion assay. See Materials and Methods. All strains contain *xylA*::*sspB^EC^* and P*amj*-*tomato*: *yumC*-ssrA*, JLG6506; *fer*-ssrA* Δ*ykuP*, JLG7141; *dnaN*-ssrA*, JLG6514. (E) Titration of viable cells in cultures of strains lacking one or both flavodoxins and in which Fer is degraded (YkuN or YkuP; “+” and “–” indicate presence and absence, respectively). All strains contain *xylA*::*sspB^EC^* and all of the Fer degradation strains contain *fer-ssrA\**. Ten-fold dilution series were spotted on LB plates without xylose or with 1% xylose. (F) RIP of strains lacking flavodoxins and in which Fer is degraded. Cultures were grown in LB medium, treated with 1% xylose to trigger degradation of Fer-ssrA*, and imaged on agarose pads during exponential phase. Scale bar, 1 μm; all panels are at the same scale. All strains contain *xylA*::*sspB^EC^*: control, JLG364; *fer-ssrA*,* JLG4330; *fer*-*ssrA\** Δ*ykuN*, JLG7070; *fer*-*ssrA\** Δ*ykuP*, JLG7072; *fer*-*ssrA\** Δ*ykuNOP*, JLG7122.

Degradation of YumC during exponential growth gave rise to elongated cells with multiple regularly-spaced chromosomes (Fig. 2C). Many cells stained with the nucleic acid stain SYTOX Green, signaling a breach of the cell’s permeability barrier. This profile matched that of cells in which the biosynthesis of the cytoplasmic PG precursors was disrupted, either by degradation of biosynthetic enzymes such as MurAA, MurC, or RacE, or by treatment with the antibiotics D-cycloserine or fosfomycin (Fig. 2B-C) (33), indicating that YumC is required for cell wall biosynthesis. When YumC was degraded in osmoprotective medium, cells lost their rod shape and developed bulges (Fig. S4). The YumC-like protein TrxB2 in *L. lactis* is required for the activity of class Ib ribonucleotide reductase in an aerobic environment (26), but the *B. subtilis* YumC profile did not match that associated with degradation of proteins such as DnaN and NrdF that are required for this function (Fig. 2B) (34), and supplementation of the media with deoxyribonucleosides did not eliminate the requirement for YumC (26; Fig. S1 and Fig. S7). Unlike *L. lactis* TrxB2, YumC appeared to be required for PG precursor biosynthesis.

Disruption of PG biosynthesis leads to induction of the σ^M^-mediated cell-wall stress response (36). We found that degradation of YumC did so as well. We fused the σ^M^-dependent promoter P*amj* (37, 38) to *tomato*, integrated the construct at *abn2*, and assessed activation on agar plates. When a filter paper disk containing xylose was placed at the center of a lawn of the YumC degradation strain, a band of Tomato fluorescence appeared at the boundary between the inhibition and growth zones, indicating activation of σ^M^ when YumC is degraded (Fig. 2D). No activation was observed when we degraded proteins that are not involved in PG biosynthesis or upon treatment with antibiotics targeting other cellular processes (Fig 2D; Fig. S5).

The requirement for YumC for PG biosynthesis would be expected to implicate one or more of the ECPs, the ferredoxin Fer or the flavodoxins YkuN or YkuP. If so, degradation of these proteins should give rise to a cytological profile associated with inhibition of PG biosynthesis and activate the σ^M^-dependent cell-wall stress response. We used RIP to obtain the cytological profile of a strain in which both flavodoxins were absent and in which Fer could be degraded. We deleted the *ykuNOP* operon and replaced *fer* with *fer-ssrA\**, so that Fer could be degraded upon induction of the resident *sspB^EC^* gene. Growth was compromised severely when we supplied the inducer xylose (Fig. 2E). The cytological profile, in which many cells were elongated with multiple chromosomes or simply lysed, resembled that which we observed when we degraded YumC, implicating these ECPs in PG biosynthesis (Fig. 2F). In order to differentiate the contributions of the two flavodoxins, we assessed viability when Fer was degraded in conjunction with a deletion of only *ykuP* or *ykuN* as opposed to a deletion of the entire operon. Growth was compromised in the *ykuP* deletion strain, but not in the *ykuN* deletion strain (Fig. 2E). RIP analysis of the *ykuP* deletion strain again pointed to PG biosynthesis (Fig. 2F), and the σ^M^ response was induced (Fig. 2D). Thus, YumC is essential for PG biosynthesis in *B. subtilis* and can make use of the ferredoxin Fer and the flavodoxin YkuP.

### Degradation of YumC disrupts the MEP isoprenoid biosynthetic pathway

Which of the many reactions of PG biosynthesis might require electrons from an FNR such as YumC and its associated ECPs? Of the reactions that generate the disaccharide pentapeptide subunit that is incorporated into the growing PG chain, only the reduction of UDP-N-acetylglucosamine-enolpyruvate (UDP-GlcNac-EP) to UDP-N-acetylmuramic acid (UDP-MurNac) requires NADPH (39, 40) (Fig. 3A). But MurB, which catalyzes this reaction, binds NADPH directly, transfers electrons from NADPH to its own FAD prosthetic group, which in turn reduces UDP-GlcNac-EP, a mechanism that obviates the requirement for exogenous electron donors such as ferredoxin or the flavodoxins (41). YumC, Fer, and YkuP should therefore not be required in this reaction.

**Fig. 3.**
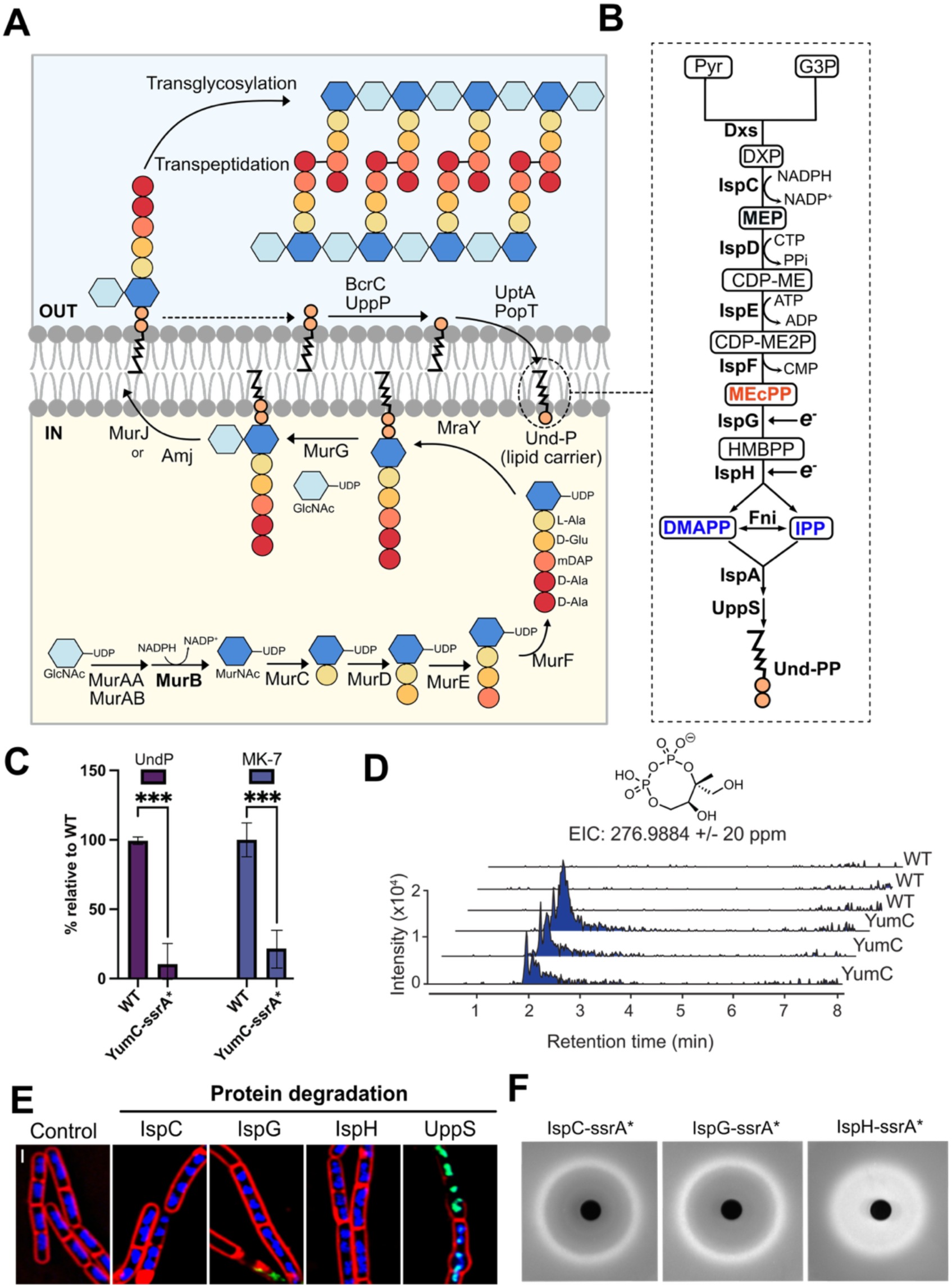
YumC is required for the biosynthesis of undecaprenyl phosphate (Und-P). (A) The peptidoglycan building block, which is assembled in the cytoplasm (yellow field), must be translocated to the outer face of the membrane (grey field) for incorporation into the growing cell wall. The building block consists of the glycan moiety N-acetylglucosamine–N-acetylmuramic acid (hexagons) amide linked to the pentapeptide L-Ala–D-Glu–meso-diaminopimelic acid–D-Ala–D-Ala (large spheres). Attachment of the lipid carrier Und-P (black squiggle) enables the building block to be flipped across the cytoplasmic membrane by the flippase MurJ or Amj (38, 68). Once the peptidoglycan building block is incorporated into the cell wall, Und-PP is released into the outer leaflet of the membrane, dephosphorylated to Und-P by BcrC or UppP (69, 70), and flipped back to the inner leaflet by UptA or PopT (71), where it can be reused. (B) Und-PP is an isoprenoid and is produced from pyruvate and glyceraldehyde 3-phosphate via the MEP pathway, two of whose enzymes, IspG and IspH, require the input of electrons from ECPs. Intermediates and products of the MEP pathway: DXP, 1-deoxy-D-xylulose 5-phosphate; MEP, 2-C-methyl-D-erythritol 4-phosphate; CDP-ME, 4-diphosphocytidyl-2-C-methyl-D-erythritol; CDP-ME2P, 4-diphosphocytidyl-2-C-methyl-D-erythritol 2-phosphate; MEcPP, 2-C-methyl-D-erythritol 2,4-cyclodiphosphate; HMBPP, 1-hydroxy-2-methyl-2-(E)-butenyl 4-diphosphate; IPP, isopentenyl diphosphate; DMAPP, dimethylallyl diphosphate. Enzymes of the MEP pathway: Dxs, DXP synthase; IspC, DXP reductoisomerase; IspD, MEP cytidylyltransferase; IspE, CDP-ME kinase; IspF, MEcPP synthase; IspG, HMBPP synthase; IspH, HMBPP reductase; Fni, IPP isomerase; IspA, farnesyl diphosphate synthase; UppS, undecaprenyl diphosphate synthase. IPP and DMAPP (blue) are the precursors for all isoprenoids. MEcPP (red) accumulates upon YumC degradation. (C) Relative undecaprenyl (Und-P, purple) and menaquinone-7 (MK-7, blue) abundance in strains containing YumC-ssrA* (JLG1728), after 2.5 h of growth in the presence of 0.5% xylose. Undecaprenyl and MK-7 levels were normalized to pellet mass and then scaled to those in the wild type (WT) strain grown in the absence of xylose. Data represent the average and standard deviation of 3 biological replicates. Significance was determined by a 2-way ANOVA with Šídák’s multiple comparisons test to determine significance between induction times; *** = p<0.001. (D) Liquid chromatography–high-resolution mass spectrometry analysis of methanol extracts from wild-type (WT) and YumC-degradation strains (YumC). Profiles from three independent biological replicates are shown. An MEcPP peak is observed in the YumC samples in each replicate. The molecular structure of MEcPP is shown above the chromatographic traces, together with the m/z value used for extracted ion chromatogram (EIC) analysis. (E) RIP of strains in which essential proteins in the MEP isoprenoid biosynthetic pathway are degraded. Cultures were grown in LB medium, treated with 1% xylose to trigger degradation of the ssrA*-tagged protein (indicated on top of each panel), and imaged on agarose pads during exponential phase. Membranes were stained with FM4-64 (red), and DNA was stained with DAPI (blue) and with SYTOX Green (green), which is ordinarily cell-impermeant. Scale bar, 1 μm; all panels are at the same scale. All strains contain *xylA*::*sspB^EC^*: control, JLG364; *ispC*-*ssrA\**, JLG5947; *ispG*-*ssrA\**, JLG6053; *ispH*-*ssrA\**, JLG1819; *uppS*-*ssrA\**, JLG6032. (F) Induction of the σ^M^ cell wall stress response upon degradation of different target proteins, assessed by disk diffusion assay. See Materials and Methods. All strains contain *xylA*::*sspB^EC^* and P*amj*-*tomato*: *ispC*-ssrA*, JLG6562; *ispG*-ssrA*, JLG6563; *ispH*-ssrA*, JLG6512.

Once all but GlcNAc is incorporated into the subunit, the monosaccharide pentapeptide is conjugated to a lipid carrier that will enable the subunit, after GlcNac is added in the next and final step, to be flipped across the cytoplasmic membrane for incorporation into the growing PG chain (Fig. 3A). The lipid carrier is undecaprenyl phosphate (Und-P), which is an isoprenoid, a derivative of isoprene. Isoprenoids are produced from the isomeric precursors isopentenyl pyrophosphate (IPP) and dimethylallyl pyrophosphate (DMAPP), which are synthesized in *B. subtilis* from glyceraldehyde 3-phosphate and pyruvate via the 2-C-methyl-D-erythritol-4-phosphate (MEP) pathway (Fig. 3B). This pathway contains three redox reactions, and all are essential (Fig. 3B; (17)). IspC, which catalyzes the first of these reactions, acquires the electrons for reduction directly from NADPH. Neither IspG nor IspH binds NADPH. Each contains iron sulfur clusters that provide electrons to reduce their substrates (recently reviewed in (42)). Biochemical experiments with these enzymes from various organisms point to a requirement for a flavodoxin or ferredoxin to supply electrons to these iron-sulfur clusters (43–45). *In vivo* data are consistent with these findings: isoprenoid biosynthesis in *E. coli* requires Flavodoxin I, one of its two flavodoxins (46).

The requirement for ferredoxin- or flavodoxin-dependent electron transfer in the MEP pathway might therefore explain why degradation of YumC or of its associated ECPs disrupts PG biosynthesis. Degradation of YumC would be expected to diminish electron flow from these carriers to their downstream targets, including the reducing enzymes of the MEP pathway, which should depress cellular isoprenoids levels. When we surveyed the metabolites present in cells in which YumC was degraded, we found that Und-P species were indeed underrepresented (Fig. 3C). Menaquinone-7 (MK-7), the only quinone that *B. subtilis* uses to shuttle electrons and whose lipid tail is heptaprenyl, a shorter chain variant of Und-P (47, 48), was also underrepresented (Fig. 3C). The substrate for IspG, 2-C-methyl-D-erythritol 2,4-cyclopyrophosphate (MEcPP) accumulated, which we determined by matching observed to predicted MEcPP fragmentation products (Fig. 3D; Fig. S6). This finding is consistent with YumC being required for the enzymatic activity of IspG. An independent experiment that entailed a scaled-up extraction with improved recovery confirmed the accumulation of MEcPP (Supplementary Methods). Because of this block at IspG, it was not possible to determine whether YumC is also required for the activity of IspH, which catalyzes the next reaction in the MEP pathway. The scaled-up extraction revealed no other metabolites with profiles that were altered consistently in response to degradation of YumC.

RIP analysis of IspC, IspG, IspH, and also of UppS, the enzyme that catalyzes the synthesis of Und-PP from IPP and farnesyl pyrophosphate, gave rise to cytological profiles resembling those from RIP analysis of YumC and of Fer in a Δ*ykuP* background (Fig. 3E; Fig. 2C and F), namely disrupted PG biosynthesis. Degradation of these proteins also induced the σ^M^ response (Fig. 3F).

YumC is thus essential in *B. subtilis* at the very least because it is required for the biosynthesis of PG. YumC is required specifically for the biosynthesis of the isoprenoid lipid carrier Und-P that enables PG building block translocation across the cytoplasmic membrane. Und-P is produced through the MEP isoprenoid biosynthesis pathway. The MEP pathway contains an enzyme (and likely two) that requires reducing equivalents from NADPH, and the transfer of these reducing equivalents is catalyzed by YumC jointly with either the ferredoxin Fer or the flavodoxin YkuP.

### Replacing the MEP pathway with the mevalonate pathway eliminates the requirement for Fer and YkuP, but not the requirement for YumC

Having implicated the FNR YumC, and the ECPs Fer and YkuP in PG biosynthesis through isoprenoid and Und-P biosynthesis, we then asked whether these proteins are also required for one or more essential functions beyond isoprenoid biosynthesis. Would *B. subtilis* tolerate the degradation of YumC or the simultaneous absence of Fer and YkuP if isoprenoid biosynthesis through the MEP pathway were rendered dispensable because IPP and DMAPP could be obtained from a difference source? To address this question, we took advantage of the fact that isoprenoids are synthesized through two independently-evolved pathways (49, 50). Most eubacteria use the MEP pathway that is present in *B. subtilis* (Fig. 3B). But archaea, animals, and fungi use the mevalonate (MVA) pathway (Fig. 4A), which makes use of an entirely different set of reactions to generate the same end-products, IPP and DMAPP. The MVA pathway is also found in some eubacteria, including *L. lactis* (51). Because the MVA pathway generates IPP and DMAPP through reactions that do not involve redox steps such as those catalyzed by IspG and IspH, it should not require an FNR such as YumC or any associated ECPs. We established the MVA pathway in *B. subtilis* and asked whether its expression would eliminate the lethality associated with degradation of YumC or degradation of Fer in a Δ*ykuP* background (Fig. 4). We engineered a synthetic operon containing the genes for the last four enzymes of the *L. lactis* MVA pathway, which catalyze the production of IPP from the metabolic intermediate mevalonate. We included the IPP isomerase that catalyzes the conversion of IPP into DMAPP, which was previously shown to be necessary for growth of MEP-pathway-deficient *E. coli* strains engineered to express the MVA pathway (52), and also the downstream *ispA* gene for farnesyl diphosphate synthase, which catalyzes the production of farnesyl diphosphate from IPP and DMAPP (Fig 4B). The synthetic operon, which we placed under the IPTG-inducible hyper-spank promoter and integrated into the chromosome at *amyE*, was functional in *B. subtilis*. When cells were supplied with mevalonate, expression of the operon rescued growth of a strain in which IspC was degraded and therefore unable to produce IPP and DMAPP through the endogenous MEP pathway (Fig. 4C). We then tested whether expression of the MVA-pathway would eliminate the lethality associated with degradation of YumC, or with degradation of Fer in a Δ*ykuP* background. Degradation of Fer abolished growth in the strain lacking YkuP, but growth was restored through expression of the MVA operon. The rescue resembled that observed in the IspC degradation strain, confirming that Fer or YkuP is required for isoprenoid biosynthesis via the MEP pathway, and revealing that this requirement is the reason why one of these ECPs must be present in *B. subtilis*. In stark contrast, expression of the MVA pathway did not restore growth to a strain in which YumC was degraded (Fig. 4C).

**Fig. 4.**
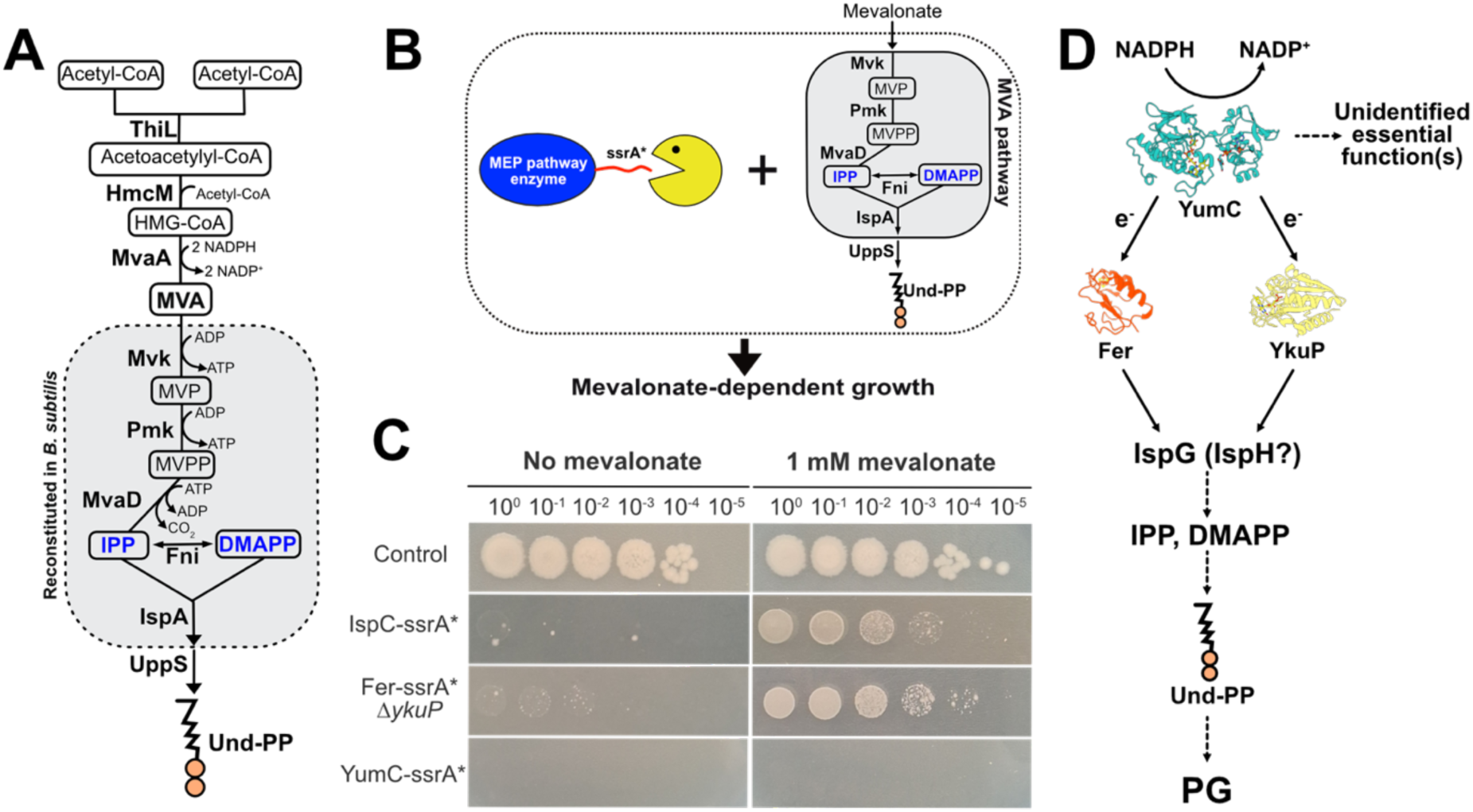
The MVA pathway complements degradation of Fer in a strain that lacks YkuP, but does not complement degradation of YumC. (A) The MVA pathway for isoprenoid biosynthesis entails the synthesis of mevalonate from three molecules of acetyl-CoA (outside the grey box), and its subsequent conversion into the isoprenoid precursors IPP and DMAPP in a series of non-redox reactions (within the grey box). The pathway from *L. lactis* is diagrammed. Enzymes: ThiL, acetoacetyl-CoA thiolase; HmcM, HMG-CoA synthase; MveA, HMG-CoA reductase; Mvk, mevalonate kinase; Pmk, phosphomevalonate kinase; MvaD, mevalonate pyrophosphate decarboxylase; Fni, IPP isomerase; IspA, farnesyl pyrophosphate synthase; UppS, undecaprenyl pyrophosphate synthase. Intermediates and products: HMG-CoA, 3-hydroxy-3-methylglutaryl coenzyme A; MVA, mevalonate; MVP, mevalonate 5-phosphate; MVPP, mevalonate pyrophosphate; IPP, isopentenyl pyrophosphate; DMAPP, dimethylallyl pyrophosphate. The enzymes within the grey box were heterologously expressed in *B. subtilis*. (B) Scheme for complementation with the MVA pathway. Degradation of enzymes of the MEP pathway (blue oval; degradation indicated by yellow pacman) is lethal. Viability can be restored by introduction of the MVA pathway. In this case, only the latter half of the mevalonate pathway was introduced (grey box). An IPTG-inducible synthetic operon encoding Mvk, Pmk, MvaD, Fni and IspA, was inserted at *amyE.* When the MEP pathway is disrupted and the MVA synthetic operon is expressed, viability would be expected to depend on supplementation with mevalonate, which is converted into IPP and DMAPP by the enzymes of the MVA-pathway. (C) Titration of viable cells in cultures of *B. subtilis* strains expressing the *L. lactis* MVA pathway. All strains contained *xylA*::*sspB^EC^* and the IPTG-inducible synthetic MVA operon. Plates contained 0.2% xylose to trigger degradation of the corresponding ssrA*-tagged protein, and 500 μM IPTG to induce the expression of MVA pathway. The series on the left was imaged from plates that also contained 1 mM mevalonate; the series on the right was imaged from plates that contained no mevalonate. Strains were as follows: control, JLG6038; *ispC-ssrA\**, JLG6042; *fer-ssrA* ΔykuP*, JLG7140; *yumC-ssrA\**, JLG6041. (D) Model for the contribution of YumC to PG biosynthesis. YumC catalyzes the transfer of electrons from NADPH to the enzymes IspG (and perhaps to IspH as well) through the ferredoxin Fer and the flavodoxin YkuP, thereby enabling the biosynthesis of IPP and DMAPP. Und-PP, which is essential for PG biosynthesis, is produced from IPP and DMAPP. YumC is also required for one or more additional essential functions, likely in a Fer- and YkuP-independent manner.

That the MVA pathway could not restore growth to a strain in which YumC was degraded indicated that YumC is required for at least one other essential process, one which apparently does not require Fer and YkuP. In *L. lactis*, the aerobic growth defect that results from the absence of the YumC-like protein TrxB2 could be offset by supplementation with deoxynucleosides. But we found that the same was not true for *B. subtilis*. Nucleoside supplementation in conjunction with expression of the MVA pathway did not restore growth when YumC was degraded (Fig. S7A). And an alternative strategy, in which we fed esterified variants of IPP and DMAPP to the cells in order to obviate the requirement for a functioning isoprenoid biosynthesis pathway, gave the same result. Neither IPP nor DMAPP can be taken up by *B. subtilis*, but esterification of the terminal phosphate to short chains of lipophilic benzoic acid derivatives eliminates their negative charges and yields cell-permeant variants of the isomeric pair (53, 54). Once in the cytoplasm, these esters are hydrolyzed by endogenous esterases, and IPP and DMAPP are released. Supplying esterified IPP to *B. subtilis* in which IspC was degraded did allow the cells to bypass isoprenoid biosynthesis, but supplying esterified IPP to *B. subtilis* in which YumC was degraded did not enable the strain to grow (Fig. S7B), and this was also the case when deoxynucleosides were supplied (Fig S7C).

## DISCUSSION

*yumC* is one of very few essential genes in *B. subtilis* whose essential function was unknown. We have demonstrated that YumC is required for the biosynthesis of isoprenoids, whose deficiency prevents PG biosynthesis (Fig. 4D). The isoprenoid Und-P is the lipid carrier that enables the PG building block to be translocated across the cytoplasmic membrane and incorporated into the cell wall. Und-P is produced through the MEP isoprenoid pathway, which contains two redox enzymes, IspG and IspH. We have demonstrated that YumC channels electrons from NADPH to IspG (and likely also IspH) through the ECPs Fer or YkuP. With this understanding of how reducing power is provided to the latter steps of the pathway, we likely have a complete inventory of the enzymes required for isoprenoid biosynthesis in *B. subtilis*.

It follows that any process that depends on isoprenoids would require YumC. In addition to serving as the lipid carrier for the PG building block, the isoprenoid Und-P is also the carrier for wall teichoic acids (55). The menaquinone MK-7 is the only quinone electron carrier in *B. subtilis,* and it contains an isoprenoid tail that is embedded in the cytoplasmic membrane (48, 56). MK-7 shuttles electrons from cytoplasmic dehydrogenases to the electron transport chain in the membrane, and is critical for respiration. *B. subtilis* also produces C35 polycyclic terpenoids that influence the properties of the spore envelope (57) and may increase spore resistance to reactive oxygen species (58). YumC would be expected to contribute to all these functions.

YumC is also likely required for a variety of processes in addition to isoprenoid biosynthesis. Biochemical and *in vivo* studies have implicated YumC-associated ECPs in biotin, nitrous oxide, and quenosine biosynthesis, and in lipid desaturation and fatty acid oxidation. Each pathway contains at least one enzyme that requires reducing electrons for its function, but do not bind NADPH directly (18–20, 22–25). And there are likely others. Any enzyme that is reduced by ferredoxin or by YkuN or YkuP would be a candidate for having a YumC requirement.

YumC from *B. subtilis* and TrxB2 from *L. lactis* are 39% identical in amino acid sequence, and the proteins are functionally interchangeable and can substitute for each other *in vivo* (26). TrxB2 is required only for aerobic growth. This is because it is required for ribonucleotide reduction, specifically for the reduction of the ribonucleotide reductase-associated flavodoxin NrdI (26). *L. lactis* has two systems for ribonucleotide reduction. One is a class III oxygen-sensitive system that can function only in an anaerobic environment, and the other, which includes NrdI, is a class Ib system that requires oxygen and can function (ostensibly) only in an aerobic environment (26, 59). YumC is essential in *B. subtilis* for both aerobic and anaerobic growth, which is unsurprising given that YumC is required for isoprenoid and cell wall biosynthesis. Because *L. lactis* makes use of the alternative MVA pathway for isoprenoid biosynthesis, TrxB2 is not required in this context. This raises the question of whether YumC is also required for ribonucleotide reduction in *B. subtilis*. *B. subtilis* has only one ribonucleotide reductase, it is a class Ib system, and it is required in both aerobic and anaerobic environments (60). The *B. subtilis* NdrI would be expected to require YumC for its reduction, as does *L. lactis* and very likely *B. cereus* as well (28), and this would amount to another essential function for YumC. But we were unable to demonstrate this experimentally. In our RIP analysis, degradation of YumC gave rise to the characteristic PG biosynthesis inhibition profile of elongated cells containing multiple regularly spaced chromosomes, from which we could also infer that DNA replication was still ongoing and that the supply of dNTPs was therefore adequate. In a strain in which DnaN or NrdF is degraded, by contrast, the elongated cells contained irregularly spaced amorphous masses of DNA (Fig. 2B) (34). We also found that YumC remained indispensable even when we introduced the MVA pathway (or supplied esterified versions of IPP), and also added exogenous deoxyribonucleosides. We infer that YumC is not likely to be required for the reduction of NdrI in *B. subtilis*, but further work is required to settle this definitively.

To perform its essential role in isoprenoid biosynthesis, YumC uses the *B. subtilis* ferredoxin, Fer, and the flavodoxin YkuP. The *ykuNOP* operon is induced when iron is limiting (61, 62), so Fer may predominate until iron grows scarce, whereupon YkuP substitutes. YkuN makes at most a minor contribution to isoprenoid biosynthesis under our experimental conditions. Degradation of Fer in a strain lacking *ykuP* impairs growth as strongly as does degradation of Fer in a strain lacking the entire *ykuNOP* operon. *In vitro* studies have shown that YumC can reduce both Fer (9) and the flavodoxin YkuN(20), but comparable evidence is lacking for YkuP. However, biochemical studies suggest that this is not an unreasonable possibility. An *E. coli* FNR can reduce both YkuN and YkuP, and either flavodoxin can reduce the heme in *B. subtilis* nitric-oxide synthase (19), transfer electrons to the cytochrome P450 BioI (22), or reduce the iron-sulfur cluster in the *B. subtilis* 7-carboxy-7-deazaguanine synthase (QueE) (25). Now we provide *in vivo* evidence that YumC can also reduce YkuP.

Our findings, taken together with the biochemical literature, raise the question of how interchangeable the two *B. subtilis* flavodoxins are. Although YkuN and YkuP are of similar size, are over 40% identical in amino acid sequence, and AlphaFold predicts that the proteins share a similar set of secondary structural elements in a similar disposition (63), no crystal structure exists for either protein. An extensive biophysical analysis found many fundamental similarities but also some notable differences between the two proteins (22). YkuN and YkuP bind FMN avidly and show nearly identical reduction potentials, but circular dichroism spectra indicate that while the structure of YkuN is unaltered with FMN bound, YkuP undergoes substantial changes to its secondary structural profile and is actually destabilized when it binds FMN. YkuP is overall less thermodynamically stable than YkuN. The behaviour of these proteins in the cell cannot of course be inferred from their biophysical properties, but it is possible, for example, that effective substrates of YkuP confer added stability upon the protein. Both biochemical and physiological studies point to selectivity, although not as stark a selectivity as we encountered with the MEP isoprenoid biosynthesis pathway. *In vitro*, it is YkuN that is more efficient in the reductive activation of nitric-oxide synthase, P450 BioI, and QueE (19, 22, 25). In isoprenoid biosynthesis *in vivo*, YkuP is the predominant flavodoxin.

We were surprised to discover that the scope of YumC’s involvement in *B. subtilis* physiology does not match precisely that of Fer and the flavodoxins. The establishment of the MVA pathway eliminated the requirement for Fer and YkuP, but not the requirement for YumC. YumC is required for at least one other essential function, but Fer and YkuP are essential only for isoprenoid synthesis. Whatever other essential function or functions YumC carries out cannot rely exclusively on Fer and YkuP. Yet, the function could still be mediated redundantly by Fer, YkuP and YkuN. In this model, any of the three ECPs could provide electrons to the same essential process—or to multiple processes whose combined disruption is lethal—distinct from isoprenoid synthesis, such that retaining only one ECP would be enough to carry out that process. It is also possible that the additional essential function of YumC is mediated by another electron carrier. The SubtiWiki data base lists 13 other proteins that contain flavodoxin domains, but of these only NrdI is essential (63). The simple solution to this puzzle is that YumC does catalyze the transfer of electrons from NADPH to NrdI for ribonucleotide reduction and that this is its additional essential function. As discussed above, this does not seem to be the case, and our inability to complement by supplying deoxyribonucleosides when the MVA pathway is active would mean that there is yet another essential function for YumC. YumC might catalyze the delivery of electrons from NADPH directly to a protein substrate, with no intervening ECP, which would be an unlikely mechanism for an FNR (7). YumC might also catalyze the delivery of electrons from NADPH directly to a small molecule, a mechanism for which there are many examples in the biochemical literature. The electron acceptor in these studies is typically a dye that loses or changes color when reduced. Even simple ionic salts such as potassium ferricyanide can serve as electron acceptors (9, 64). This diaphorase activity has been exploited in biochemistry studies for purification and characterization of FNRs, including YumC (9). It remains to be seen whether this *in vitro* activity of the YumC protein has any implications *in vivo* for *B. subtilis*.

Although YumC and TrxB2 are functionally interchangeable, the essential roles that they perform are distinct and unrelated, owing to the difference in the metabolic configuration of their host organisms. This suggests that equivalent FNR/ECP redox modules can transduce reducing potential from NADPH to a variety of redox enzymes in a variety of different organisms. This versatility may enable the FNR/ECP modules to contribute to the many metabolic pathways for which they are required. The versatility arises from the ability of the FNR to interact with both classes of ECPs, the ferredoxins and flavodoxins (7), and more remarkably, from the ability of these ECPs to interact with a wide variety of redox enzymes. If a pathway to which the FNR/ECP module contributes is essential, as is the case for isoprenoid biosynthesis in *B. subtilis*, then the module is itself essential. It follows that uncovering the reason why the module is essential in a one organism may give little to no insight as to why it is essential in another organism. This, and the fact that the module will typically contribute to several pathways within a single organism, explains why physiological studies of YumC and the ECPs of *B. subtilis* have lagged behind biochemical studies. In the present study, RIP was able to lead us to an essential function of YumC, its requirement in PG biosynthesis, and we could then follow this lead back to isoprenoid biosynthesis and to IspG. The requirement for *B. subtilis* YumC and ferredoxin or YkuP to provide reducing power to this redox enzyme is so far the only unambiguous explanation for YumC being essential in *B. subtilis*.

## MATERIAL AND METHODS

### Strains and plasmids

Strains are listed in Table S1, plasmids in Table S2, and oligonucleotides in Table S3. All *B. subtilis* strains used in this study are derivatives of *Bacillus subtilis* PY79. Strain and plasmid constructions are described in the Supplementary Information.

### Cre-mediated antibiotic cassette excision

Plasmid pJLG5875 was constructed to facilitate the Cre-mediated excision of antibiotic resistance genes flanked by *loxP* sites, which was frequently required in the course of strain construction. The *cre* gene was placed under control of a xylose-dependent promoter and cloned into a mini variant of the *B. subtilis natto* plasmid pLS20 (65), with its segregation system engineered to promote rapid loss of the plasmid in the absence of antibiotic selection (66).

Strains containing an antibiotic resistance gene flanked by *lox* sites were transformed with pJLG5875, with selection for tetracycline resistance (10 μg/mL). Transformants were purified once on a tetracycline plate, then passaged onto a plate containing 0.5% xylose, and the plate was incubated overnight. Colonies that arose on the xylose plate were passaged onto an LB plate, and the plate was incubated overnight. The LB passage was repeated, and colonies that arose after the second LB passage were screened for loss of resistance to the target antibiotic and to tetracycline. Loss of both antibiotic resistances, indicating successful Cre-mediated excision and subsequent loss of pJLG5875, was typically in the range of 80 to 100%. Although we typically performed the procedure at 30 °C, it was equally effective at 37 °C.

### Rapid Inhibition Profiling (RIP)

RIP was performed as described (33). Briefly, a preculture was started by inoculating 5 mL of liquid LB medium with a single colony from an LB plate that had been incubated overnight at 30 °C. The preculture was grown at 37 °C in a roller drum to an OD_600_ of 0.2-0.5, diluted 1:100 into 5 mL of LB, and then incubated at 37 °C in a roller drum until OD_600_ 0.11-0.15. At this point, xylose was added to the culture to a final concentration of 1% in order to induce the degradation of the ssrA*-tagged protein. Samples were taken for microscopy at hourly intervals for up to three hours. Cells were visualized on 1% agarose pads prepared with Ultrapure agarose (Invitrogen, cat#16500500) in LB diluted 1/4. The pads were supplemented with FM4-64 at 1 mg/mL (Invitrogen, cat#T3166), SYTOX Green at 0.25 mM (Invitrogen, cat#S7020) and DAPI at 200 μg/mL (Carl ROTH, cat#6335.1). Eight μl of culture were transferred to the agarose pad. Phase contrast and fluorescence images were acquired with a DeltaVision Ultra Microscope equipped with a PCO Edge sCMOS camera. Imaging conditions were as follows: FM4-64, excitation 542 nm/emission 679 nm/exposure time 0.200 s/light transmission 20%; SYTOX Green, excitation 475 nm/emission 525 nm/exposure time 0.005 s/light transmission 2%; DAPI excitation 390 nm/emission 435 nm/exposure time 0.100 s/light transmission 40%. Images were deconvolved with SoftWoRx v5.5.1 (Applied Precision). Z-stacks were processed with Fiji (67). Medial focal planes are shown.

To examine cell morphology in osmoprotective medium (Fig. S4), cells were grown in LB-MSM agarose pads containing 1% xylose to trigger degradation of ssrA*-tagged proteins and the pads imaged after 3 h of incubation at 30 °C. LB-MSM was prepared by combining one part 2X magnesium-sucrose-maleic acid (MSM) buffer (40 mM MgCl₂, 1 M sucrose, and 40 mM maleic acid, pH 7) with one part 2X LB (33).

### Anaerobic growth of *B. subtilis*

*B. subtilis* is a facultative anaerobe that can use nitrate as an electron acceptor (31). LB agar plates containing 0.5% KNO_3_ and 0.5% arabinose were poured in aerobic conditions and pre-incubated overnight in an anaerobic chamber (5% H_2_, 5% CO_2_, and 90% N_2_). Plates also contained 0.2% xylose or both 0.2% xylose and 4 mM deoxynucleosides as required for the experiment of Fig. S2. Strains were streaked from cryostocks onto LB plates and grown aerobically overnight at 30 °C. The plates were then transferred to the anaerobic chamber and colonies were streaked onto the pre-incubated plates. The plates were incubated in the anaerobic chamber for 2 days.

### Disk diffusion assays

LB cultures were grown at 30 °C in a roller drum to an OD_600_ of 0.4-0.6. 200 µL of the culture was added to 4 mL of top agar (0.7% agar), and the mixture poured atop a standard 1.5% agar LB plate. 6-mm diameter Whatmann Antibiotic Assays Discs (Cytiva Whatmann, cat#2017-006) were applied to the plates and spotted with 5 µL of antibiotic (10 mg/mL) or xylose (20%). The plates were incubated overnight at 30 °C and photographed the next day with the ChemiDoc Imaging System (Bio-Rad) using an excitation light source around 530 nm (Green Epi) and an emission filter between 602 and 650 nm. All images were captured with an exposure time of 1.0 s.

### Mevalonate and deoxynucleoside complementation assays

LB cultures were grown at 37 °C with aeration to an OD_600_ slightly above 1. The OD_600_ of the cultures were normalized to 1, diluted serially 10-fold, and 5 μL of each dilution were spotted onto LB agar supplemented with 0.5 mM IPTG (Thermo Scientific, cat#R0392), 0.2% xylose (Sigma-Aldrich, cat#X1500) and 1 mM DL-mevalonate (Thermo Scientific Chemicals, cat#428300050) prepared as previously described by Martin *et al*. (52).

For supplementation with deoxynucleosides, a mixture of deoxyadenosine (Sigma-Aldrich, cat#D8668), deoxyguanosine (Sigma-Aldrich, cat#854999), deoxycytidine (Sigma-Aldrich, cat#D0776), and thymidine (Sigma-Aldrich, cat#T1895), was used at a concentration of 4 mM (1 mM each) for agar plates and 0.4 mM (0.1 mM) for liquid cultures. Plates were incubated at 37 °C overnight and photographed the next day.

### Culture and extraction conditions for targeted metabolomics

A 17 mm × 100 mm culture tube (VWR, Radnor, PA) was prepared with 5 mL of LB broth. An exponentially growing *B. subtilis* culture at OD_600_ of 1 was added to each 5 mL culture at a 1:20 dilution. Cultures were incubated in a shaking incubator (New Brunswick Scientific, Edison, NJ) at 37 °C and 250 rpm for 3 h, then induced by addition of xylose to a final concentration of 0.5% and returned to the incubator. At 1 h post-induction, half of the cultures (n = 3) were transferred to 15 mL conical tubes (VWR, Radnor, PA) and centrifuged at 4,000 × *g* for 6 min. The supernatants were discarded, and the cell pellets were flash-frozen in liquid nitrogen and lyophilized overnight using a benchtop freeze-dryer (Labconco, Kansas City, MO). The remaining cultures (n = 3) were incubated for an additional 1.5 h (2.5 h total post-induction) and harvested as described above.

The following day, dry pellet masses were recorded, and pellets were extracted in 10 mL of methanol. Samples were bath-sonicated (Branson Ultrasonics, Danbury, CT) at room temperature for 30 min and then rotated on a tube rotator (Fisher Scientific, Waltham, MA) at 25 rpm for 2 h. Samples were centrifuged at 4,000 × *g* for 6 min, and the supernatants were transferred to new 15 mL conical tubes and dried under nitrogen using a Reacti-Vap evaporator (Thermo Fisher, Waltham, MA). Dried extracts were reconstituted in 100 μl of isopropanol and transferred to 1.5 mL microcentrifuge tubes. Extracts were clarified by centrifugation at 10,000 × *g* for 5 min, and the supernatants were transferred to LC/MS vials (Thermo Fisher, Waltham, MA).

## ACKNOWLEDGMENTS

We thank members of the Evolutionary Cell Biology group and the Microbial Population Biology department of the Max Planck Institute for Evolutionary Biology for continuous discussion and feedback. We acknowledge the National BioResource Project (NIG, Japan): *B. subtilis* for shipping the BKE collection to us. This work was supported by ERC starting grant 853323, the National Science Foundation Chemistry of Life Processes (NSF-CLP) program (CHE-2204170), and the National Institute of General Medical Sciences at the National Institutes of Health (1R35GM157211-01).

## AUTHOR CONTRIBUTIONS

Conceptualization, DA, AID, JLG; Methodology, DA, MCA, DPM, AID, JAB, JLG; Validation, DA, MCA, DPM, AID; Formal analysis, DA, MCA, DPM, AID; Investigation, DA, MCA, DPM, AID; Resources, AL; Data Curation, DA, MCA, DPM, AID; Writing - Original Draft, MCA, AID, JLG; Writing - Review & Editing, All authors; Writing - Fina Draft, AID, JLG; Visualization, MCA, AID, JLG; Supervision, AID, JAB, JLG; Project administration, JLG; Funding acquisition, JAB, JLG.

