## Supplementary Figures and Tables for "The ferredoxin/flavodoxin-NADP^+^ oxidoreductase YumC is essential for isoprenoid and peptidoglycan biosynthesis in *Bacillus subtilis*"

**Contents:**

- Supplementary Figures 1 to 7
- Supplementary Tables 1 to 3
- Supplementary Methods:
  - Molecular biology
  - Plasmid construction
  - Genetic selections for mutations that bypass YumC essentiality
  - Scaled-up *B. subtilis* extraction for broader *YumC* metabolite survey
  - Liquid chromatography high-resolution mass spectrometry (LC-HRMS) quantification by ESI
  - Liquid chromatography high-resolution mass spectrometry with data-dependent acquisition (LC-HRMS DDA) by ESI
  - Liquid chromatography high-resolution mass spectrometry (LC-HRMS) quantification of MK-7 and undecaprenyl species by APCI
  - Growth Curve Analyses with *B. subtilis*

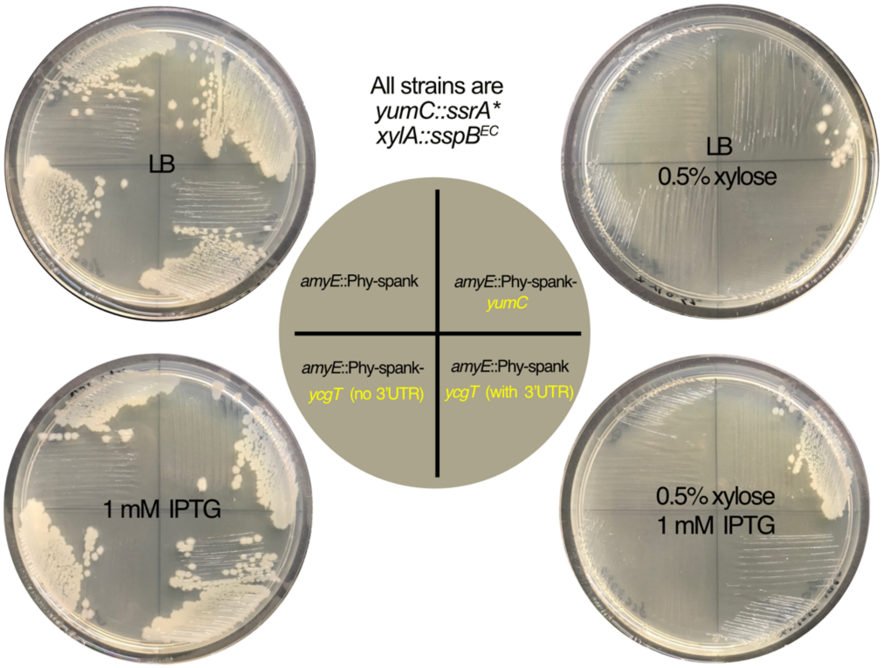


**Figure S1. Expression of *ycgT* does not complement YumC-ssrA* degradation.** All strains are derivatives of the YumC-degradation strain JLG1728 (PY79 *yumC-ssrA**, *xylA::sspB*$^{EC}$). Colonies were streaked from LB agar plates onto four plates according to the schema in the center of the figure (upper left quadrant, JLG3767; upper right quadrant, JLG3948; lower right quadrant, JLG3950; lower left quadrant, JLG3949). The LB agar plates were supplemented as indicated in the figure, with 0.5% xylose to induce degradation of YumC, with 1 mM IPTG to induce expression of an ectopic copy of *yumC* or *ycgT*. Induction of YumC degradation prevented growth of all four strains (upper right plate). Coordinate induction of ectopic *yumC*, but not of ectopic *ycgT*, complemented YumC degradation (lower right plate). The few colonies that arose on the upper right plate are due most likely to leaky expression of *yumC* from $P_{hyper\text{-}spank}$ or to easily arising mutations in loci that are required for YumC degradation (see Figure S3, below.)

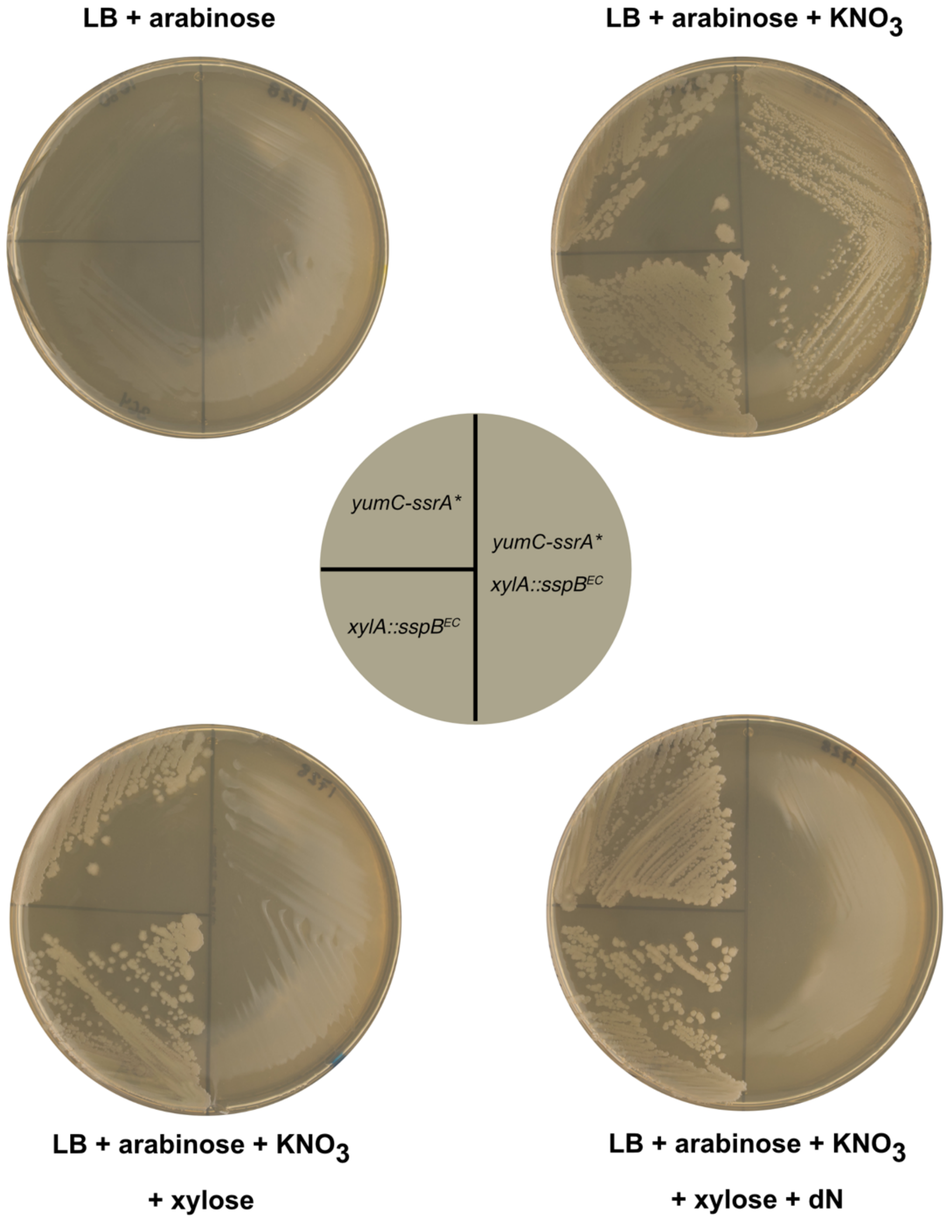


**Figure S2. YumC is essential for anaerobic growth.** Colonies were streaked from LB agar plates onto four plates according to the schema in the center of the figure (upper left quadrant, JLG1680; lower left quadrant, JLG364; right half, JLG1728). The LB agar plates were supplemented as indicated in the figure, with arabinose at 0.5%, with $KNO_3$ at 0.5%, with deoxyribonucleosides (dN) at 1 mM each, and with xylose at 0.2% to induce degradation of YumC. Plates were incubated anaerobically for 48 h at 37°C. Anaerobic growth requires the presence of $NO_3$, confirming that the strains are carrying out anaerobic respiration (upper plates), YumC is required for anaerobic growth (righthand plates), and dN do not compensate for the absence of YumC during anaerobic growth (lower right plate).

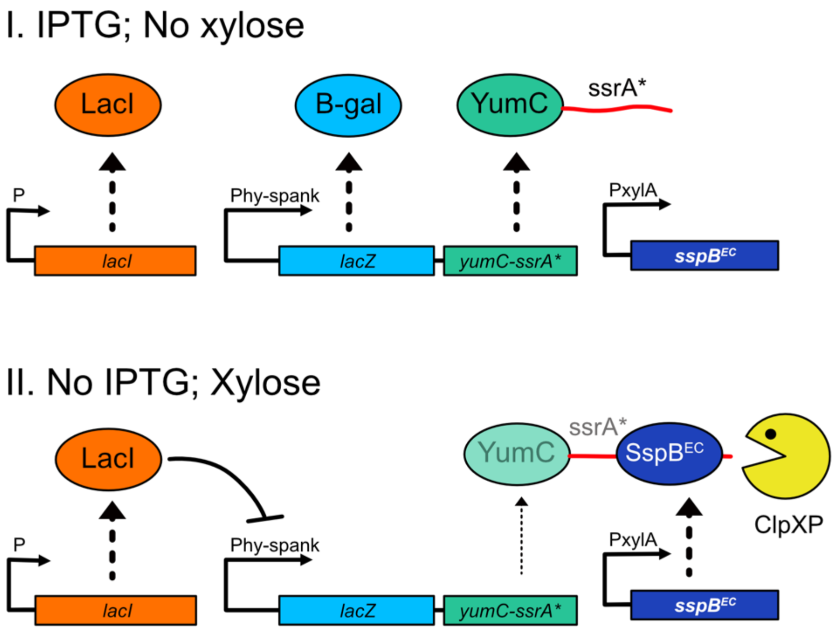


**Figure S3. Genetic selection for mutations that suppress the lethality of YumC depletion**. (A) Design of the selection. A synthetic operon consisting of *lacZ* (light blue box) and *yumC-ssrA** (green box) under the control of the IPTG-inducible hyper-spank promoter (Phy-spank) was inserted into the chromosome at *amyE*. The native copy of *yumC* was deleted, so that *yumC-ssrA** was the only source of YumC. The chromosome also contained *lacI* expressed constitutively (orange box) and *sspB*$^{EC}$ inserted at *xylA* and under the control of the xylose inducible *xylA* promoter (dark blue box). In the presence of IPTG and the absence of xylose, such that the synthetic operon is expressed and both β-galactosidase (light blue oval) and YumC-ssrA* (green oval) are produced (upper diagram), the strain forms blue colonies on agar plates containing X-Gal. In the absence of IPTG and the presence of xylose (lower diagram), the synthetic operon is not expressed. Any YumC that is produced nevertheless would be recognized by SspB$^{EC}$ (dark blue oval) and targeted for degradation by ClpXP (yellow Pacman). With YumC absent, the strain is inviable and no colonies arise. Mutants were selected that were able to grow in the absence of IPTG and the presence of xylose. Mutants that produced blue colonies on X-Gal plates were not investigated as they likely contained mutations that simply enabled the synthetic operon to be expressed in the absence of IPTG. Of the approximately $10^{11}$ cells in total on which the selection was performed, only 12 white colonies were recovered. (B) Whole genome sequencing of the candidates. All of the mutants that gave rise to white colonies contained a dramatic expansion of the *yumC-ssrA** region, with reads 10- to 30-fold more abundant than the rest of the chromosome (blue bars). The expansion did not include the *lacZ* gene, and likely originated via recombination between a pair of 46 bp direct repeats, one upstream of *yumC-ssrA**, the other downstream of *lacI*. This increase in *yumC-ssrA** copy number presumably resulted in steady state levels of YumC that were high enough to compensate for both the transcriptional repression and YumC-ssrA* degradation. Instead of bypassing the requirement for YumC, these mutations bypassed the rigorous selection constraints and enabled enough YumC to be produced for the mutant to grow.

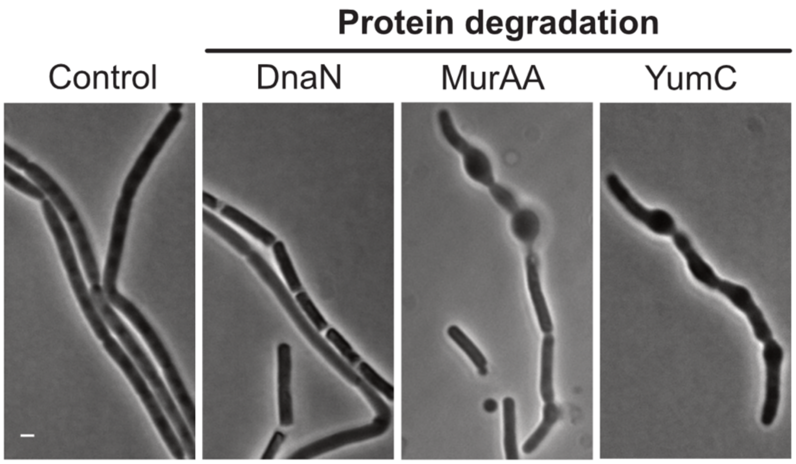


**Figure S4. YumC-depleted cells deform in osmoprotective medium.** Phase contrast images of cells in which selected proteins are degraded in osmoprotective medium. Cells were grown on LB-MSM agarose pads for 3 hours at 30°C. Pads contained 1% of xylose to trigger degradation of the ssrA*-tagged protein indicated at the top of each panel. Scale bar, 1 µm; all panels are at the same scale. All strains contain *xylA*::*sspB*$^{EC}$: control, JLG364; *dnaN-ssrA**, JLG715; *murAA-ssrA**, JLG736; *yumC-ssrA**, JLG1728.

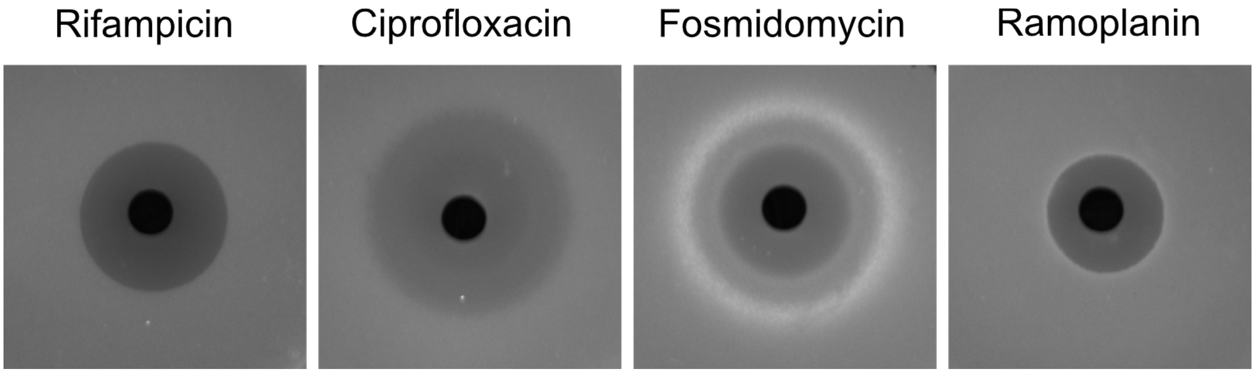


**Figure S5. Activation of the $\sigma^M$-mediated cell-wall stress response in the presence of various antibiotics.** Induction of the $\sigma^M$ cell wall stress response in *B. subtilis* PY79 assessed by disk diffusion assay. Rifampicin inhibits transcription by binding to the β subunit of RNA polymerase. Ciprofloxacin inhibits DNA gyrase. Fosmidomycin inhibits the enzyme IspC from the MEP pathway for isoprenoid synthesis. Ramoplanin inhibits cell wall synthesis by binding and sequestering lipid II. All the antibiotics were at a concentration of 10 μg/ml.

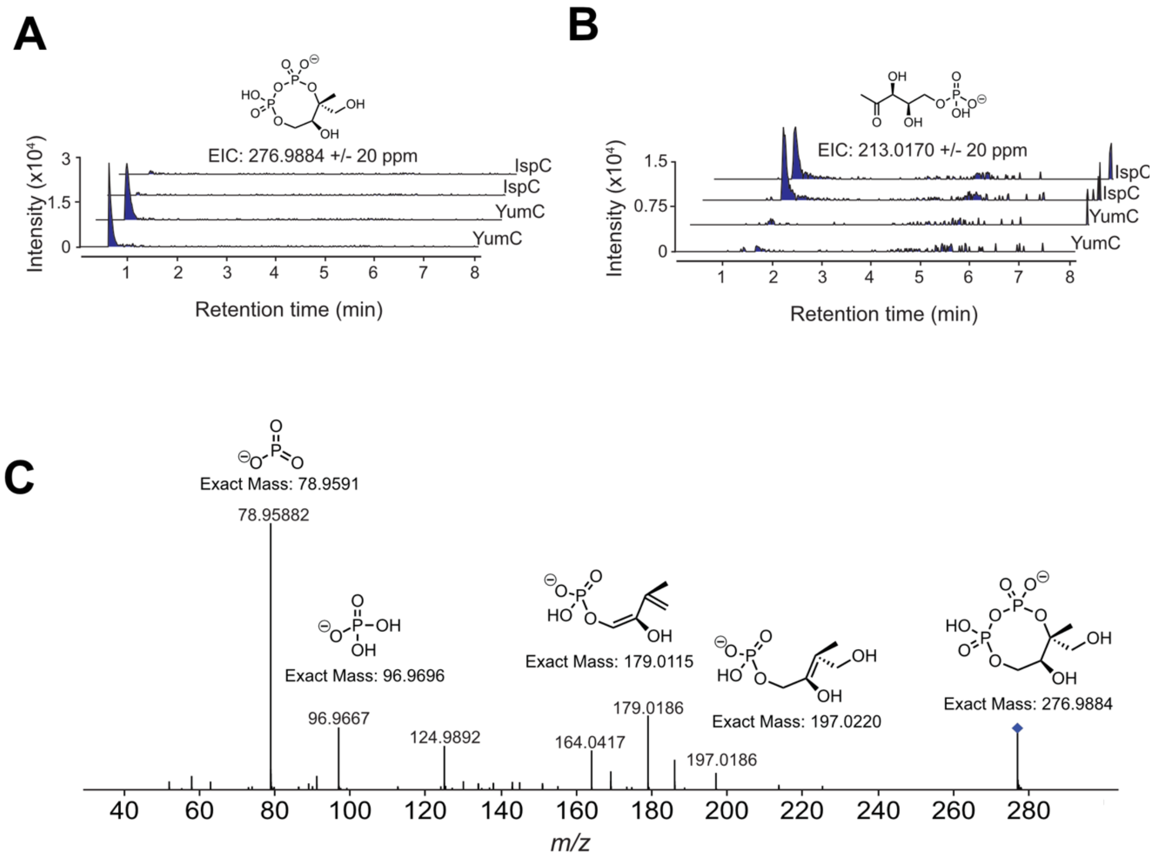


**Figure S6**. **Degradation of YumC and of IspC leads to the accumulation of distinct MEP pathway intermediates.** Liquid chromatography–high-resolution mass spectrometry analysis of methanol extracts from the the IspC degradation strain JLG5947 (IspC), and the YumC degradation strain JLG1728 (YumC) following xylose-induced degradation. Profiles from two independent biological replicates are shown. (A) An MEcPP peak is present in the YumC-degradation samples but absent from the IspC-degradation samples. The molecular structure of the IspG substrate MEcPP is depicted above the chromatographic traces, together with the m/z value used for extracted ion chromatogram (EIC) analysis. (B) A DXP peak is observed in the IspC-degradation samples but is absent from the YumC-degradation samples. The molecular structure of the IspC substrate DXP is depicted above the chromatographic traces, together with the m/z value used for extracted ion chromatogram (EIC) analysis. (C) MS/MS fragmentation profile for MEcPP.

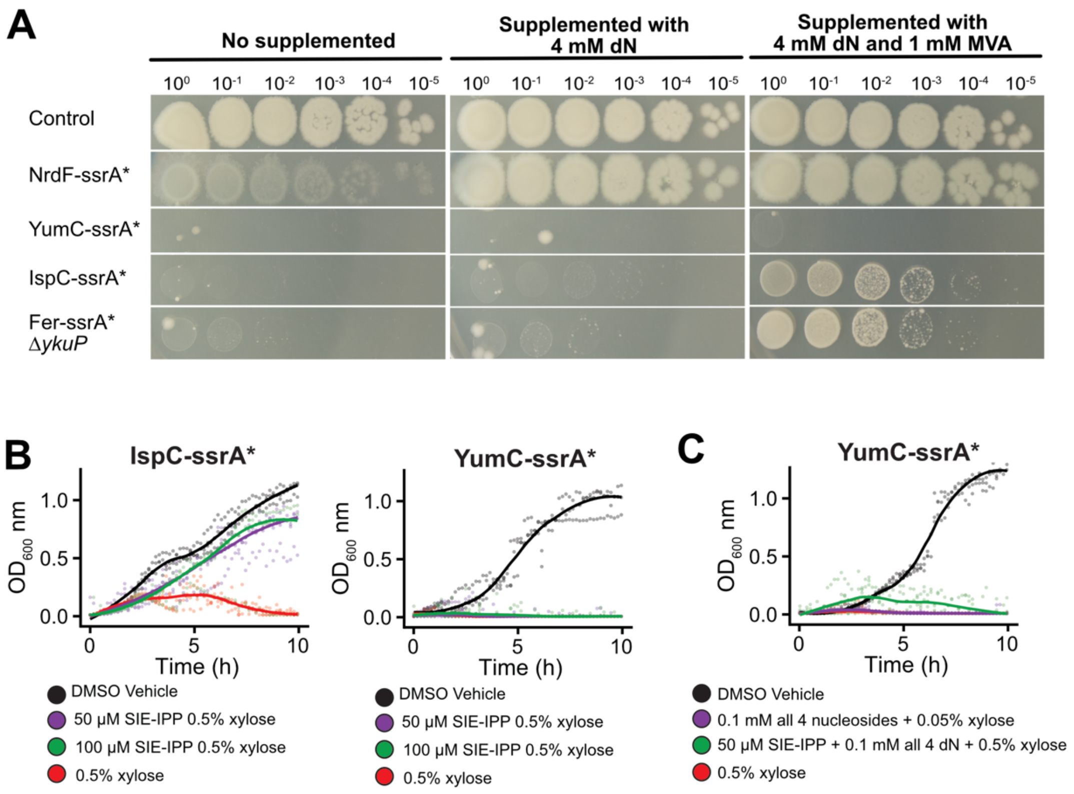


**Figure S7. Media supplementation that rescues IspC degradation or Fer degradation in the absence of YkuP does not rescue YumC degradation** (A) Titration of viable cells in cultures of strains supplemented with deoxyribonucleosides and in which the mevalonate synthetic operon is activated. Plates contained 0.2% xylose to induce production of $SspB^{EC}$ and degradation of the ssrA*-tagged protein, and 0.5 mM IPTG to induce the expression of the mevalonate (MVA) synthetic operon. The lefthand panels correspond to plates that were not supplemented, the middle panels to plates supplemented with 4 mM deoxynucleosides (dN; 1mM each), and the righthand panels to plates supplemented with 1 mM MVA and 4 mM dN. All strains except NrdF-ssrA are derivatives of strain JLG6038, which contains the MVA synthetic operon integrated into the chromosome at *amyE*: Control, JLG6038; NrdF-ssrA*, JLG1251; YumC-ssrA*, JLG6041; IspC-ssrA*, JLG6042; Fer-ssrA* Δ*ykuP*, JLG7140. Activation of the MVA pathway with or without dN supplementation (see Fig. 4C) rescues both degradation of IspC and degradation of Fer in the absence of YkuNOP, but not degradation of YumC. (B) Growth curves of IspC and YumC degradation strains supplemented with SIE-IPP, an esterified derivative of IPP that is cell-permeant. Growth curves of IspC (left) and YumC (right) degradation strains in LB supplemented with DMSO (black), with 0.5% xylose to induce the degradation of the ssrA*-tagged proteins (red), with 0.5% xylose and 50 μM SIE-IPP (purple), with 0.5% xylose and 100 μM SIE-IPP (green). SIE-IPP supplementation rescues degradation of IspC but not of YumC. (C) Growth curves of the YumC degradation strain in LB supplemented with DMSO (black), with 0.5% xylose to induce the degradation of the ssrA*-tagged proteins (red), with 0.5% xylose and 0.4 mM dN (purple), or with 0.5% xylose and 50 μM SIE-IPP and 0.4 mM dN (green). None of these supplementations rescues degradation of YumC. Strains are IspC-ssrA*, JLG5947; YumC-ssrA*, JLG1728.

| **Table S1. Strains used in this study** | | |
|---|---|---|
| **Strain** | **Genotype or description** | **Reference, source or construction[a]** |
| *B. subtilis* 168 | *trpC2* | ATCC 23857 |
| *B. subtilis* PY79 | Wild type | (1) |
| BKE14150 | *trpC2* Δ*ykuN*::Em | (2) |
| BKE14170 | *trpC2* Δ*ykuP*::Em | (2) |
| BKK23040 | *trpC2* Δ*fer*::Em | (2) |
| JLG241 | *murC-sfGFP-ssrA**ΩKm | pJLG43 → PY79 |
| JLG364 | *ΔxylA*::*sspB*$^{EC}$ΩCm | (3) |
| JLG380 | *murC-sfGFP-ssrAΩKm* Δ*xylA*::*sspB*$^{EC}$ΩCm | JLG241 → JLG364 |
| JLG707 | *fabZ-ssrA**ΩKm Δ*xylA*::*sspB*$^{EC}$ΩCm | (3) |
| JLG715 | *dnaN-ssrA**ΩKm Δ*xylA*::*sspB*$^{EC}$ΩCm | (3) |
| JLG736 | *murAA-ssrA**ΩKm Δ*xylA*::*sspB*$^{EC}$ΩCm | (3) |
| JLG1170 | *tagO-sfGFP-ssrA**ΩKm Δ*xylA*::*sspB*$^{EC}$ΩCm | (4) |
| JLG1245 | *racE-ssrA**ΩKm Δ*xylA*::*sspB*$^{EC}$ΩCm | (3) |
| JLG1251 | *nrdF-ssrA**ΩKm Δ*xylA*::*sspB*$^{EC}$ΩCm | (4) |
| JLG1680 | *yumC-ssrA**ΩKm | pJLG254 → PY79 |
| JLG1728 | *yumC-ssrA**ΩKm Δ*xylA*::*sspB*$^{EC}$ΩCm | JLG1680 → PY79 (Km) |
| JLG1819 | *ispH-ssrA*ΩKm* Δ*xylA*::*sspB*$^{EC}$ΩCm | JLG3122 → JLG364 (Km) |
| JLG2319 | *rpoC-ssrA**ΩKm | pER172 → PY79 (Km) |
| JLG2320 | *rpsB-ssrA**ΩKm | (5) |
| JLG3122 | *ispH-ssrA**ΩKm | pER68 → PY79 (Km) |
| JLG3306 | *accA-ssrA**ΩKm | (3) |
| JLG3767 | *yumC-ssrA**ΩKm Δ*xylA*::*sspB*$^{EC}$ΩCm *amyE*::$P_{hyperspank}$ΩSp | pDR111 → JLG1728 (Sp) |
| JLG3782 | Δ*fer*::*loxP*-Km-*loxP* | BKK23040 → PY79 (Km) |
| JLG3822 | *ispG-ssrA**ΩKm | pJLG769 → PY79 (Km) |
| JLG3833 | *fer-ssrA**ΩKm | pJLG794 → PY79 (Km) |
| JLG3948 | *yumC-ssrA**ΩKm Δ*xylA*::*sspB*$^{EC}$ΩCm *amyE*::$P_{hyperspank}$-*yumC*ΩSp | JLG4020 → JLG1728 (Sp) |
| JLG3949 | *yumC-ssrA**ΩKm Δ*xylA*::*sspB*$^{EC}$ΩCm *amyE*::$P_{hyperspank}$-*ycgT* without long 3'UTR S120ΩSp | JLG4030 → JLG1728 (Sp) |

| | | |
|---|---|---|
| JLG3950 | yumC-ssrA*ΩKm ΔxylA::$sspB^{EC}$ΩCm amyE::$P_{hyperspank}$-ycgT with long 3'UTR S120ΩSp | JLG4066 → JLG1728 (Sp) |
| JLG4020 | amyE::$P_{hyperspank}$-yumCΩSp | pJLG7274 → PY79 (Sp) |
| JLG4030 | amyE::$P_{hyperspank}$-ycgT without long 3'UTR S120ΩSp | pJLG7275 → PY79 (Sp) |
| JLG4066 | amyE::$P_{hyperspank}$-ycgT with long 3'UTR S120ΩSp | pJLG7276 → PY79 (Sp) |
| JLG4330 | fer-ssrA*ΩKm ΔxylA::$sspB^{EC}$ΩCm | JLG3833 → JLG364 (Km) |
| JLG5154 | ΔykuNOP::Em | pJLG1224 → PY79 (MLS) |
| JLG5946 | ispC-ssrA*ΩKm | pJLG1239 → PY79 (Km) |
| JLG5947 | ispC-ssrA*ΩKm ΔxylA::$sspB^{EC}$ΩCm | JLG5946 → JLG364 (Km) |
| JLG6020 | amyE::$P_{hyperspank}$-lacZ-yumC-ssrA*ΩSp | pJLG1232 → PY79 |
| JLG6021 | amyE::$P_{hyperspank}$-lacZ-yumC-ssrA*ΩSp ΔxylA::$sspB^{EC}$ΩCm | JLG6020 → JLG364 |
| JLG6023 | amyE::$P_{hyperspank}$-lacZ-yumC-ssrA*ΩSp ΔxylA::$sspB^{Ec}$ΩCm ΔyumC::Em | pJLG1229 → JLG6021 |
| JLG6031 | uppS-ssrA*ΩKm | pJLG1249 → PY79 (Km) |
| JLG6032 | uppS-ssrA*ΩKm ΔxylA::$sspB^{EC}$ΩCm | JLG6031 → JLG364 (Km) |
| JLG6038 | amyE::$P_{hyperspank}$-mvaD-pmk-mvk-fni-ispAΩSp | pJLG1236 → PY79 (Sp) |
| JLG6041 | yumC-ssrA*ΩKm ΔxylA::$sspB^{EC}$ΩCm amyE::$P_{hyperspank}$-mvaD-pmk-mvk-fni-ispAΩSp | JLG6038 → JLG1728 (Sp) |
| JLG6042 | ispC-ssrA*ΩKm ΔxylA::$sspB^{EC}$ΩCm amyE::$P_{hyperspank}$-mvaD-pmk-mvk-fni-ispAΩSp | JLG6038 → JLG5947 (Sp) |
| JLG6053 | ispG-ssrA*ΩKm ΔxylA::$sspB^{EC}$ΩCm | JLG3822 → JLG364 (Km) |
| JLG6506 | yumC-ssrA*ΩKm ΔxylA::$sspB^{EC}$ΩCm abnB2::$P_{amj}$-tomatoΩEm | JLG6511 → JLG1728 (MLS) |
| JLG6511 | abnB2::$P_{amj}$-tomatoΩEm | pJLG1342 → PY79 (MLS) |
| JLG6512 | ispH-ssrA*ΩKm ΔxylA::$sspB^{EC}$ΩCm abnB2::$P_{amj}$-tomatoΩEm | JLG6511 → JLG1819 (MLS) |
| JLG6514 | dnaN-ssrA*ΩKm ΔxylA::$sspB^{EC}$ΩCm abnB2::$P_{amj}$-tomatoΩEm | JLG6511 → JLG715 (MLS) |
| JLG6549 | ΔykuN::Em | BKE14150 → PY79 (MLS) |
| JLG6550 | ΔykuP::Km | BKE14170 → PY79 (Km) |
| JLG6562 | ispC-ssrA*ΩKm ΔxylA::$sspB^{EC}$ΩCm abnB2::$P_{amj}$-tomatoΩEm | JLG6511 → JLG5947 (MLS) |

| | | |
|---|---|---|
| JLG6563 | *ispG-ssrA**ΩKm Δ*xylA*::*sspB*$^{EC}$ΩCm *abnB2*::*P*$_{amj}$-*tomato*ΩEm | JLG6511 → JLG6053 (MLS) |
| JLG6654 | Δ*ykuN* | pJLG5875 → JLG6549 (Tet) |
| JLG6655 | Δ*ykuP* | pJLG5875 → JLG6550 (Tet) |
| JLG7066 | *fer-ssrA**ΩKm Δ*ykuN* | JLG4330 → JLG6654 (Km) |
| JLG7068 | *fer-ssrA**ΩKm Δ*ykuP* | JLG4330 → JLG6655 (Km) |
| JLG7070 | *fer-ssrA**Ωkm Δ*ykuN* Δ*xylA*::*sspB*$^{EC}$ΩCm | JLG364 → JLG7066 (Cm) |
| JLG7072 | *fer-ssrA**ΩKm Δ*ykuP* Δ*xylA*::*sspB*$^{EC}$ΩCm | JLG364 → JLG7068 (Cm) |
| JLG7122 | *fer-ssrA**ΩKm Δ*xylA*::*sspB*$^{EC}$ΩCm Δ*ykuNOP*ΩEm | JLG5154 → JLG4330 (MLS) |
| JLG7140 | *fer-ssrA**ΩKm Δ*ykuP* Δ*xylA*::*sspB*$^{EC}$ΩCm *amyE*::*P*$_{hyperspank}$-*mvaD-pmk-mvk-fni-ispA*ΩSp | JLG6038 → JLG7072 (Sp) |
| JLG7141 | *fer-ssrA**ΩKm Δ*ykuP* Δ*xylA*::*sspB*$^{EC}$ΩCm *abnB2*::*P*$_{amj}$-*tomato*ΩEm | JLG6511→JLG7072 (MLS) |
| JLG7234 | *rpsB-ssrA**ΩKm Δ*xylA*::*sspB*$^{EC}$ΩCm | JLG2320→JLG364 (Cm) |
| JLG7235 | *rpoC-ssrA**ΩKm Δ*xylA*::*sspB*$^{EC}$ΩCm | JLG2319→JLG364 (Cm) |

[a]Plasmid or genomic DNA (left side of the arrow) used to transform an existing strain (right side of the arrow) to construct a new strain is listed. The antibiotic used for selection is given in parentheses (Km, kanamycin; MLS, erythromycin and lincomycin; Cm, chloramphenicol; Sp, spectinomycin). Antibiotic concentrations were as follows: kanamycin, 50 μg/ml (for *E. coli* TOP10) and 10 μg/ml (for *B. subtilis*); chloramphenicol, 5 μg/ml; MLS, 1 μg/mL erythromycin and 25 μg/mL lincomycin; spectinomycin, 100 μg/ml.

| Table S2. Plasmids used in this study | |
|---|---|
| **Plasmid** | **Description/Reference** |
| pAID3244 | (6) |
| pBR329 | (7) |
| pCrePA | (8) |
| pWH1520 | (9) |
| pDR111 | Gift from David Rudner |
| pER68 | pDG1662(Sp-ori-Ap)-*ispH-ssrA**ΩKm |
| pER172 | pDG1662(Sp-ori-Ap)-*rpoC-ssrA**ΩKm |
| pJLG3 | (10) |
| pJLG36 | (10) |
| pJLG43 | pBR329(ori)-*murC-sfGFP-ssrA**ΩKm |
| pDG1731 | (11) |
| pDG1662 | (11) |
| pJLG254 | pDG1662(Sp-ori-Ap)-*yumC-ssrA**ΩKm |
| pJLG769 | pDG1662(Sp-ori-Ap)-*ispG-ssrA**ΩKm |
| pJLG794 | pDG1662(Sp-ori-Ap)-*fer-ssrA**ΩKm |
| pJLG884 | (12) |
| pJLG1060 | pBR329(ori)-*yxiA::Pcv1-tomato*ΩEm |
| pJLG1224 | pDG1662(Sp-ori-Ap)-Δ*ykuNOP*::Em |
| pJLG1229 | pDG1662(Sp-ori-Ap)-Δ*yumC*::*loxP*-Em-*loxP* |
| pJLG1230 | pDR111-*lacZ-yumC*ΩSp |
| pJLG1231 | pDR111-$RBS_{tufA}$-*lacZ-yumC*ΩSp |
| pJLG1232 | pDR111-$RBS_{tufA}$-*lacZ-yumC-ssrA**ΩSp |
| pJLG1233 | pDR111-*mvaD-pmk*ΩSp |
| pJLG1234 | pDR111-*mvaD-pmk-mvk*ΩSp |
| pJLG1236 | pDR111-*mvaD-pmk-mvk-fni-ispA*ΩSp |
| pJLG1239 | pDG1662(Sp-ori-Ap)-*IspC-ssrA**ΩKm |
| pJLG1249 | pDG1662(Sp-ori-Ap)-*UppS-ssrA**ΩKm |
| pJLG1342 | pBR329(ori)-*yxiA::Pamj-tomato*ΩEm |
| pJLG5875 | pAID3244-*cre* (Tet) |
| pJLG7274 | pDR111-*yumC*ΩSp |
| pJLG7275 | pDR111-*ycgT*ΩSp |
| pJLG7276 | pDR111-*ycgT*-3'UTRΩSp |

| **Table S3. Oligonucleotides used in this study** | |
|---|---|
| **Primer** | **Sequence**[a] |
| oER227 | gggttaacgcgtaatccatgGATATCACGTGATCTATATCGGC |
| oER228 | catcatttgctgcgctagcGTTTTTTGCTTTTACTTTTGGAAG |
| oER229 | cactggagttgtcccaattGGCTGTAAAAGCCTCAGTTTTTTTATAG |
| oER230 | cacatttccccgaaaagtgcGAGCTTGTTCATGAAGTAAGGC |
| oER749 | gggttaacgcgtaatccatgCAGAAGGTTCAATCGATCCG |
| oER750 | catcatttgctgcgctagcTTCAACCGGGACCATATCG |
| oER751 | cactggagttgtcccaattCTGATTTAACTCTGCTGAAAGACTGC |
| oER752 | cacatttccccgaaaagtgcTAACCCACATACCCAGGTGG |
| oJLG7 | AATTGGGACAACTCCAGTG |
| oJLG63 | CTTCCTGGACAGGGATATGG |
| oJLG69 | GCGGCTCTTACCAGCCTAAC |
| oJLG70 | GTCTCGCGGTATCATTGCAG |
| oJLG73 | ctgcaatgataccgcgagacTCTTTCTTAATCGGAGACGG |
| oJLG74 | cttgcgcttgcgctgctagcTGCCATGACGTTTTCGTAG |
| oJLG75 | cactggagttgtcccaattCTAAAAAGCAGTGATCTCACTGC |
| oJLG76 | gttaggctggtaagagccgcGATCTGTCCAGACTGATCCG |
| oJLG77 | GCTAGCAGCGCAAGCGC |
| oJLG95 | CATGGATTACGCGTTAACCC |
| oJLG96 | GCACTTTTCGGGGAAATGTG |
| oJLG184 | GCTAGCGCAGCAAATGATG |
| oJLG253 | TCCTTTAACTCTGGCAACCC |
| oJLG254 | ATGTGATAACTCGGCGTATG |
| oJLG724 | gggttaacgcgtaatccatgAAGCGGTTGAATCTGTTGAG |
| oJLG725 | catcatttgctgcgctagcTTTATTTTCAAAAAGACTTGTTGAGTG |
| oJLG726 | cactggagttgtcccaattCTAATAAAAAAAGGAGCTTGTGTCTG |
| oJLG727 | cacatttccccgaaaagtgcAATCGTCTGCATATTGGCAG |
| oJLG1597 | TTCTGCTCCCTCGCTCAG |
| oJLG1598 | CAGGGAGCACTGGTCAAC |
| oJLG1696 | CATGGATTACGCGTTAACCCAGGTCTAGAGGATCGATCTG |
| oJLG1699 | GCTAGCAGCGCAAGCGCAGCTAGCGCAGCAAATGATG |
| oJLG1752 | gggttaacgcgtaatccatgATCAATCCGCCATATCCTGC |
| oJLG1755 | cacatttccccgaaaagtgcAATCGTCTGCATATTGGCAG |
| oJLG1760 | gggttaacgcgtaatccatgGGAAATTCAGCACTGGTGG |
| oJLG1761 | gcctgagcgagggagcagaaCCAACTCATATTCCTGAAGCG |
| oJLG1762 | gcgttgaccagtgctccctgCCAGAAGGTGAAGAAGAGG |
| oJLG1763 | cacatttccccgaaaagtgcGATCAAGCATTGGGATCGC |
| oJLG1782 | gtact*gtcgac*CATTTAGGAGGCAATTTTCGTATGC |

| | |
|---|---|
| oJLG1783 | ctgta*gcatgc*CTTCAAATAAAAAAGGAGCTGTGTC |
| oJLG1787 | ctgta*gcatgc*AAAAAAAGAGCCTAGATTAATCCAGGCT |
| oJLG1791 | ctgta*gcatgc*CTACGCGTACGGAAAAACGCTC |
| oJLG1802 | gtact*gtcgac*GATGTGAACGGGAGGCAG |
| oJLG1872 | AAGCTAATTCGGTGGAAACGAG |
| oJLG1926 | gcctgagcgagggagcagaaGCATACGAAAATTGCCTCCT |
| oJLG1927 | gcgttgaccagtgctccctgCTGCGACACAGCTCCTT |
| oJLG2091 | GACTAAGCTTAATTGTTATCCGCTC |
| oJLG2092 | GACCATTTAGGAGGCAATTTTCGTATG |
| oJLG2093 | caattaagcttagtcCAATTTCACACAGGAAACAGC |
| oJLG2094 | gcctcctaaatgGTCCTGCCCGGTTATTATTATTTTTGAC |
| oJLG2537 | 5'P-AAGGAGGATTTTAGAATGACCATGATTACGGATTCACTG |
| oJLG2538 | 5'P-AAGAGCTTTTAATTACTAAGCTTAATTGTTATCCGCTCAC |
| oJLG2572 | 5'P-TTCAGAAAATTATGCACTTGGAGGATAATAATAAAAAAAGGAGCTTGTGTCTG |
| oJLG2573 | 5'P-TAGTTTTCATCATTTGCTGCGCTAGCTTTATTTTCAAAAAGACTTGTTGAGTG |
| oJLG2657 | gcctgagcgagggagcagaaCTTCATCCTCTTCGTCTTGG |
| oJLG2658 | gcgttgaccagtgctccctgAAGTACATCCGCAACTGTCC |
| oJLG2724 | gggttaacgcgtaatccatgAGGAGGAACAATGCCG |
| oJLG2725 | tgcgcttgcgctgctagcTGTGAGTATTGAATTGACGTA |
| oJLG2726 | cactggagttgtcccaattTAAGGTGGTATGTTCGTG |
| oJLG2727 | cacatttccccgaaaagtgcCGGAAGCAGCCTGAT |
| oJLG2742 | gggttaacgcgtaatccatgCTGCAAAATCACTCGCG |
| oJLG2743 | tgcgcttgcgctgctagcAATTCCGCCAAACCTCC |
| oJLG2745 | cactggagttgtcccaattAAGGAGGCACATTAGAGGATGGTGGACATGA |
| oJLG2746 | cacatttccccgaaaagtgcAAAGACCGATCAAACCCG |
| oJLG2930 | **GAGG**AAATAACAAATGTCCAATTTACTGACCGTACACC |
| oJLG2931 | GATTAATCCAGGCTCCTAATCGCCATCTTCCAGCAGG |
| oJLG2932 | caattaagatagttgatggataaacttgttcacttaaatca**aagGAGG**AAATAACAAATGTCCAATTTACTG |
| oJLG2933 | gaatccgttagcgaggtgccgccggcttccatAAAAAAACAGCCTAGATTAATCCAGGCTCCTAATC |
| oJLG2934 | ATGGAAGCCGGCGGCAC |
| oJLG2935 | CTTTGATTTAAGTGAACAAGTTTATCCATC |
| oJLG2962 | gataacaattaagcttagtc**CTCTTAAGGAGGATTTTAGA**ATGAAAAATATTGTTACTGCAC |
| oJLG2963 | cgtttccaccgaattagcttCAGGCTCGTACAGGC |

| | |
|---|---|
| oJLG2968 | GTTTAACCTGCTAAAAACTAAAA |
| oJLG2969 | **ttgtattcctccttatatgt**GATCTCTGTCTTCAGTTATTTTTTTGAGCAATC |
| oJLG2970 | **acatataaggaggaatacaa**ATGACTATCAATAAAATGGGG |
| oJLG2971 | ttttagtttttagcaggttaaacTTAAAAGGAGTAAATCCACG |
| oJLG2996 | ctctataaataatcttatcTTAAAAGGAGTAAATCCACG |
| oJLG2997 | gataagatta**tttatagagGAGGAAATACA**ATGATGAAAAGTGAAAAAGAAATCC |
| oJLG2998 | agtaaaaacctccttctgatgctaagataTTATTTTTTTCTTTGTTGGATAAAATCG |
| oJLG2999 | tatcttagc**atcagaaggaggtttttact**ATGGATACGAAGATATTAAAGC |
| oJLG3000 | ttttagtttttagcaggttaaacTTATTCCACTTCCAGTTGTTC |
| oJLG3069 | TAAGGAGGATTTTAGAATGGTTAGC |
| oJLG3071 | ccatatccctgtccaggaagGTTTACTTTCCAGAATGATTCTTC |
| oJLG3072 | gctaaccattctaaaatcctccttaATTTCATCCTAGAGATAAGACTGG |
| 23040_Back_F | cactggagttgtcccaattTAGAAGTAACTAAAAAAAGCTCCCG |
| 23040_Back_R | cacatttccccgaaaagtgcGTAGGGCAAGTTGCGAATTT |
| 23040_Front_F | gggttaacgcgtaatccatgAAGGTGTTGTCGAAGTTCCT |
| 23040_Front_R | tgcgcttgcgctgctagcTTCAAATTTAAGCGGGTCGC |
| 25070_Back_F | cactggagttgtcccaattCGGGAACCCGTCAAGCTTTT |
| 25070_Back_R | cacatttccccgaaaagtgcGGTGGCAAACGATTACGAAA |
| 25070_Front_F | gggttaacgcgtaatccatgTTTTAAGCAAAGGCATCGGG |
| 25070_Front_R | tgcgcttgcgctgctagcAGCTTTTTGTGTTTCTTCTT |

[a]Homology regions for Gibson assembly in lower case, ribosome binding sites (RBS) in bold, and restriction sites in italics. 5'P indicates that the oligo is phosphorylated at 5'.

**SUPPLEMENTARY METHODS**

### Molecular biology

Standard techniques of molecular biology were used. Genomic DNA, plasmids and DNA fragments, including products of PCR amplifications, were purified with kits from Machery-Nagel or Qiagen. Phusion High-Fidelity DNA Polymerase and Q5 High-Fidelity DNA Polymerase from New England Biolabs (NEB) were used for PCR amplifications. Plasmids were constructed via isothermal (Gibson) assembly with the NEBuilder HiFi DNA Master Mix & Cloning Kit (NEB) or via conventional cloning with restriction endonucleases. Molecular biology reagents, including deoxynucleotide triphosphates (dNTPs), restriction endonucleases, and T4 DNA ligase, were obtained from NEB. RNase A was obtained from Qiagen. SeaKem agarose for routine gel electrophoresis was obtained from Biozym. Oligonucleotide primers (see Table S3) were synthesized by Integrated DNA Technologies (IDT). DNA sequencing was performed in house by the sequencing facility at the Max Planck Institute for Evolutionary Biology or at the Institut für Klinische Molekularbiologie at Christian-Albrechts-Universität. Plasmids were introduced into *E. coli* TOP10 by transformation of cells made competent by a variant of the Hanahan protocol. The selection was done using different antibiotics, 50 μg/mL (for *E. coli* TOP10) and 10 μg/mL (for *B. subtilis*) kanamycin, 5 μg/mL chloramphenicol, MLS (1 μg/mL erythromycin and 25 μg/mL lincomycin), 100 μg/mL spectinomycin and 100 μg/mL ampicillin.

### Plasmid construction

**pER172.** Constructed by four-piece Gibson Assembly reaction: *rpoC* upstream homology region, PCR amplified from PY79 genomic DNA with primers oER749 and oER750; ssrA*-Km fragment, PCR amplified from pJLG3 plasmid template with primers oJLG184 and oJLG7; *rpoC* downstream homology, PCR amplified from PY79 genomic DNA template with primers oER751 and oER752; and a plasmid backbone containing the origin of replication, spectinomycin resistance gene, and ampicillin resistance gene from plasmid pDG1662 (11), amplified with primers oJLG95 and oJLG96.

**pER68.** Constructed by four-piece Gibson Assembly reaction: *ispH* upstream homology region, PCR amplified from PY79 genomic DNA with primers oER227 and oER228; ssrA*-Km fragment, PCR amplified from pJLG3 plasmid template with primers oJLG184 and oJLG7; *ispH* downstream homology, PCR amplified from PY79 genomic DNA template with primers oER229 and oER230;

and a plasmid backbone containing the origin of replication, spectinomycin resistance gene, and ampicillin resistance gene from plasmid pDG1662 (11), amplified with primers oJLG95 and oJLG96.

**pJLG43.** Plasmid pJLG43 was constructed by assembling the following four fragments using Gibson assembly: (i) a 909 bp fragment consisting of the 3' region *murC* coding sequence without the stop codon, amplified from genomic DNA of *B. subtilis* PY79 with primers oJLG73 and oJLG74; (ii) a fragment containing *sfGFP-ssrA** linked to a kanamycin-resistance gene, amplified from plasmid pJLG36 with primers oJLG7 and oJLG77 from (10); (iii) a fragment consisting of the 900 pb immediately after the stop codon of *murC*, amplified from genomic DNA of *B. subtilis* PY79 with primers oJLG75 and oJLG76; (iv) a fragment containing the origin of replication from plasmid pBR329, amplified with primers oJLG69 and oJLG70.

**pJLG254.** Plasmid pJLG254 was constructed by assembling the following four fragments using Gibson assembly: (i) a fragment consisting of the last 734 bp of *yumC* coding sequence without the stop codon, amplified from genomic DNA of *B. subtilis* PY79 with primers oJLG724 and oJLG725; (ii) a fragment containing the *ssrA** tag linked to a kanamycin-resistance gene, amplified from plasmid pJLG3 with primers oJLG7 and oJLG184 (10); (iii) a 604-bp fragment corresponding to the region immediately after the stop codon of *yumC*, amplified from genomic DNA of *B. subtilis* PY79 with primers oJLG726 and oJLG727; (iv) a plasmid backbone containing the origin of replication, spectinomycin resistance gene, and ampicillin resistance gene from plasmid pDG1662 (11), amplified with primers oJLG95 and oJLG96.

**pJLG769.** Plasmid pJLG769 was constructed by assembling the following four fragments using Gibson assembly: (i) a 452-bp fragment consisting of the 3' region of *ispG* coding sequence without the stop codon, amplified with primers 25070_Front_F and 25070_Front_R (ii) a fragment containing the *ssrA** tag linked to a kanamycin-resistance gene amplified from genomic DNA of strain JLG3306 with primers oJLG7 and oJLG1699 (3); (iii) a 411-bp fragment consisting of the region immediately after the stop codon of *ispG*, amplified from genomic DNA of *B. subtilis* PY79 with primers 25070_Back_F and 25070_Back_R; (iv) a plasmid backbone containing the origin of replication, spectinomycin resistance gene, and ampicillin resistance gene from plasmid pDG1662 (11), amplified with primers oJLG96 and oJLG1696.

**pJLG794.** Plasmid pJLG794 was constructed by assembling the following four fragments using Gibson assembly: (i) the last 122 bp of the *fer* coding sequence without the stop codon, amplified from genomic DNA of *B. subtilis* PY79 with primers 23040_Front_F and 23040_Front_R, (ii) a fragment containing the *ssrA** tag linked to a kanamycin-resistance gene amplified from genomic DNA of strain JLG3306 with primers oJLG7 and oJLG1699 (3); (iii) a 387-bp fragment consisting of the region immediately after the stop codon of *fer*, amplified from genomic DNA of *B. subtilis* PY79 with primers 23040_Back_F and 23040_Back_R; (iv) a plasmid backbone containing the origin of replication, spectinomycin resistance gene, and ampicillin resistance gene from plasmid pDG1662 (11), amplified with primers oJLG96 and oJLG1696.

**pJLG1060.** Plasmid pJLG1060 was constructed by amplifying plasmid pJLG884 (12) with primers oJLG2657 and oJLG2658, and joining the amplicon with an erythromycin resistance gene amplified from plasmid pDG1731 (11) with primers oJLG1597 and oJLG1598 using Gibson assembly. The kanamycin resistance gene from plasmid pJLG884 is replaced with an erythromycin resistance gene.

**pJLG1224.** Plasmid pJLG1224 was constructed by assembling the following four fragments using Gibson assembly: (i) A 593-bp fragment containing 409 bp of the region upstream from *ykuN* and the first 184 bp of *ykuN* coding sequence, amplified from genomic DNA of *B. subtilis* PY79 with primers oJLG1760 and oJLG1761; (ii) an erythromycin resistance gene amplified from plasmid pDG1731 (11) with primers oJLG253 and oJLG254; (iii) a 438-bp fragment containing the last 135 bp of *ykuP* and the 303 bp immediately downstream of it, amplified from genomic DNA of *B. subtilis* PY79 with primers oJLG1762 and oJLG1763; (iv) a plasmid backbone containing the origin of replication, spectinomycin resistance gene, and ampicillin resistance gene from plasmid pDG1662 (11), amplified with primers oJLG96 and oJLG1696.

**pJLG1229.** Plasmid pJLG1229 was constructed by assembling the following four fragments using Gibson assembly: (i) a fragment containing the region immediately upstream of *yumC* coding sequence, amplified from genomic DNA of *B. subtilis* PY79with primers oJLG1752 and oJLG1926; (ii) an erythromycin resistance gene amplified from plasmid pDG1731 (11) with primers oJLG253 and oJLG254; (iii) a fragment containing the region immediately downstream of *yumC* coding sequence, amplified from genomic DNA of *B. subtilis* PY79 with primers oJLG1755 and oJLG1927, (iv) a plasmid backbone encompassing the origin of replication, spectinomycin resistance gene, and ampicillin resistance gene from plasmid pDG1662 (11), amplified with primers oJLG96 and oJLG1696.

**pJLG1230.** Plasmid pJLG1230 was constructed by assembling the following two fragments using Gibson assembly: (i)the *lacZ* gene from *E.coli* MG1655 genomic DNA, amplified with primers oJLG2093 and oJLG2094; (ii) an inverse PCR product of plasmid pJLG7274, amplified with primers oJLG2091 and oJLG2092.

**pJLG1231.** Plasmid pJLG1231 was constructed via inverse PCR of plasmid pJLG1230 with 5'-phosphorylated primers oJLG2537 and oJLG2538 to replace the native ribosome binding site (RBS) of *lacZ* with the RBS of *tufA* from *B. subtilis*. The amplicon was gel-purified and circularized with T4 DNA ligase.

**pJLG1232.** Plasmid pJLG1232 was constructed via inverse PCR of plasmid pJLG1231 with 5'-phosphorylated primers oJLG2572 and oJLG2573 to add the ssrA* tag to *yumC*. The amplicon was gel-purified and circularized with T4 DNA ligase.

**pJLG1233.** Plasmid pJLG1233 was constructed by assembling the following two fragments using Gibson assembly: (i) an inverse PCR product of plasmid pDR111 with primers oJLG1872 and oJLG2091; (ii) the *mvaD* and *pmk* genes, amplified from genomic DNA of *Lactococcus lactis* subsp. lactis with primers oJLG2962 and oJLG2963. oJLG2962 adds the RBS of *tufA* upstream of *mvaD*.

**pJLG1234.** Plasmid pJLG1234 was constructed by assembling the following two fragments using Gibson assembly: (i) inverse PCR fragment of plasmid pJLG1233 with primers oJLG2968 and oJLG2969 (ii) *mvk* gene, amplified from genomic DNA of *Lactococcus lactis* subsp. lactis with primers oJLG2970 and oJLG2971. oJLG2970 add the RBS of *tsf* upstream of of *mvk*.

**pJLG1236.** Plasmid pJLG1236 was constructed by assembling the following three fragments using Gibson assembly: (i) inverse PCR fragment of plasmid pJLG1234 with primers oJLG2968 and oJLG2996; (ii) *fni* gene, amplified from genomic DNA of *Lactococcus lactis* subsp. lactis with primers oJLG2997 and oJLG2998. The RBS of *alaS* from *B. subtilis* is placed upstream of *fni*; (iii) *ispA* gene, amplified from genomic DNA of *Lactococcus lactis* subsp. lactis with primers oJLG2999 and oJLG3000. The RBS of *leuS* from *B. subtilis* is placed upstream of *ispA*.

**pJLG1239.** Plasmid pJLG1239 was constructed by assembling the following four fragments using Gibson assembly: (i) a fragment consisting of the last 202 bp of the *ispC* coding sequence without the stop codon, amplified from genomic DNA of *B. subtilis* PY79 with primers oJLG2724 and oJLG2725; (ii) a fragment containing the *ssrA** tag linked to a kanamycin-resistance gene amplified from genomic DNA of strain JLG3306 with primers oJLG7 and oJLG1699 (3); (iii) a 195 bp-fragment consisting of the region immediately after the stop codon of *ispC*, amplified from genomic DNA of *B. subtilis* PY79 with primers oJLG2726 and oJLG2727; (iv) a plasmid backbone containing the origin of replication, spectinomycin resistance gene, and ampicillin resistance gene of pDG1662 (11), amplified with primers oJLG96 and oJLG1696.

**pJLG1249.** Plasmid pJLG1249 was constructed by assembling the following four fragments using Gibson assembly: (i) a fragment consisting of the last 278 bp of *uppS* coding sequence without the stop codon, amplified with primers oJLG2742 and oJLG2743 from genomic DNA of *B. subtilis* PY79; (ii) a fragment containing the *ssrA** tag linked to a kanamycin-resistance gene amplified with the primers oJLG7 and oJLG1699 from genomic DNA of strain JLG3306 (3); (iii) a fragment consisting of the 190 bp immediately after the stop codon of *uppS*, amplified with primers oJLG2745 and oJLG2746 from genomic DNA of *B. subtilis* PY79; (iv) a plasmid backbone containing the origin of replication, spectinomycin resistance gene, and ampicillin resistance gene of pDG1662 (11), amplified with primers oJLG96 and oJLG1696.

**pJLG1342**. Plasmid pJLG1342 was constructed by assembling the following two fragments Gibson assembly: (i) a fragment of 246 bp containing *amj* promoter (*Pamj*) without the ribosome binding site of *amj*, amplified from genomic DNA of *B. subtilis* PY79 with primers oJLG3071 and oJLG3072; (ii) an inverse PCR fragment of plasmid pJLG1060 amplified with primers oJLG63 and oJLG3069 to remove the *pcv1* promoter, maintaining the ribosome binding site of *tomato*. After assembly, *pcv1* promoter is replaced by *amj* promoter.

**pJLG5875**. Plasmid pJLG5875 was constructed by introducing the *cre* gene, under control of the *Bacillus megaterium* xylose promoter, into plasmid pAID3244 (6). The *cre* gene was first amplified from plasmid pCrePA (8) with primers oJLG2930 and oJLG2931, which modify the ribosome binding site of the *xylA* gene and add part of the *alp7AR* transcriptional terminator to the 3'end of the gene. The resulting amplicon was then amplified with primers oJLG2932 and oJLG2933, which completes the transcription terminator and adds flanking sequences from plasmid pWH1520 (9). This 1162 bp amplicon was then joined by Gibson assembly to a segment of pWH1520 that had been amplified with primers oJLG2934 and oJLG2935. The resulting plasmid was then restricted with NheI and NsiI, and the 7206 bp fragment ligated to the 3501 bp fragment generated by restriction of pAID3244 with NheI and NsiI.

**pJLG7274**. Plasmid pJLG7274 was constructed by amplification of genomic DNA from *B. subtilis* PY79 with primers oJLG1782 and oJLG1783, restriction of the amplicon with SalI and SphI, and ligation of the 1079 bp product to plasmid pDR111 restricted with SalI and SphI. The cloned segment contains the *yumC* ribosome binding site and the *yumC* gene through its transcriptional terminator.

**pJLG7275**. Plasmid pJLG7275 was constructed by amplification of genomic DNA from *B. subtilis* 168 with primers oJLG1802 and oJLG1787, restriction of the amplicon with SalI and SphI, and ligation of the 1078 bp product to plasmid pDR111 restricted with SalI and SphI. The cloned segment includes the *ycgT* ribosome binding site, the *ycgT* gene, and the *alp7AR* transcription terminator (13).

**pJLG7276**. Plasmid pJLG7276 was constructed by amplification of genomic DNA from *B. subtilis* 168 with primers oJLG1802 and oJLG1791, restriction of the amplicon with SalI and SphI, and ligation of the 3491 bp product to plasmid pDR111 restricted with SalI and SphI. The cloned segment contains the *ycgT* ribosome binding site, the *ycgT* gene, and its 2448 bp 3'UTR (S120).

### Genetic selections for mutations that bypass YumC essentiality

Two genetic selections were performed for mutations that suppressed the degradation of YumC-ssrA*.

*Selection 1.* A single colony of strain JLG1728, which carries *yumC-ssrA** linked to a kanamycin-resistance gene and $sspB^{EC}$ inserted in the *xylA* locus and linked to a chloramphenicol-resistance gene, was streaked across an LB agar plate containing D-(+)-xylose at 1%, 0.3%, 0.1%, 0.03%, or 0.01%, and the plate was incubated at 30°C. These concentrations of xylose prevented normal growth of JLG1728. Colonies that emerged over the next few days were restreaked onto fresh plates of the same selective medium. In total, 41 colonies were purified, and their mutations characterized: 4 from 1% xylose, 2 from 0.3% xylose, 14 from 0.1% xylose, 14 from 0.3% xylose, and 7 from 0.01% xylose. All mutants were found to carry mutations that disrupted the degradation system. Nine mutants had mutations that mapped to *yumC-ssrA** that either disrupted the ssrA* tag or prevented its addition at the C-terminus of YumC. These mutations were identified by back-crossing their YumC-ssrA* alleles to a clean background carrying *xylA*::*sspB*$^{EC}$. The resulting strains were able of grow in the presence of xylose. Sequencing of the *yumC-ssrA** region confirmed the presence of mutations. The remaining 32 mutants contained mutations that otherwise disrupted the degradation system. This was demonstrated by replacing the native *murC* with *mur*C-*ssrA** or the native *murF* with *murF-ssrA**, whose degradation should render the strain inviable, and finding instead that the resulting strain was able to grow in the presence of xylose. Of these mutants, four had mutations that were sensitive to chloramphenicol and were presumed to have mutations that eliminated *xylA*::*sspB*$^{EC}$, which was linked to a chloramphenicol-resistance marker. Five mutants were subjected to whole-genome sequencing and were found to have mutations in *clpX*: point mutations (2), in-frame deletions (2), or insertions (1). The remaining mutants were not further characterized genetically.

*Selection 2*. This selection was engineered to be more rigorous and prevent the accumulation of mutations that interfered with the degradation of YumC-ssrA*. The selection strain JLG6023 contained an IPTG-inducible synthetic operon wherein *yumC-ssrA** is preceded by the *lacZ* gene from *E. coli*. The strain also carried *xylA::sspB*$^{EC}$. The native copy of *yumC* was deleted, so that *yumC-ssrA** in the synthetic operon was the sole source of YumC. Growth required the presence of IPTG and the absence of xylose, and colonies were blue in plates containing X-Gal. Three rounds of selection were carried out, wherein 6 mL exponential cultures of JLG6023 grown in LB containing 0.5 mM IPTG were pelleted, the cells washed four times with 3 mL T-Base, and 10% and 90% of the cells plated on LB agar containing 0.5% xylose and 40 μg/mL X-Gal. In total, we screened 30 cultures, amounting to ~$10^{10}$-$10^{11}$ cells. We obtained 12 clones able to grow in the absence of IPTG and that formed white colonies in plates with X-Gal. Whole genome sequencing showed that all the clones contained an expansion of *yumC-ssrA**, with read depth 10 to 30 times higher than that of the flanking sequence. The expansion did not include the *lacZ* gene. See Figure S3 for additional details.

### Scaled-up *B. subtilis* extraction for broader *YumC* metabolite survey

A 250 mL baffled flask was prepared with 23.5 mL of LB broth. An exponentially growing *B. subtilis* culture at an $OD_{600nm}$ of 1 was added to each 25 mL culture at a 1:20 dilution. Cultures were incubated in a shaking incubator (New Brunswick Scientific, Edison, NJ) at 37 °C and 180 rpm for 2.5 h, then induced by the addition of xylose to a final concentration of 0.001% and returned to the incubator. At 2.5 h post-induction, cultures were transferred to 40 mL conical tubes (VWR, Radnor, PA) and centrifuged at 4,000 × *g* for 6 min. The supernatants were discarded, and the cell pellets were resuspended in 50 mM Tris-HCl and digested with 5 µL of a 1 mg/mL stock of T4 lysozyme (New England Biolabs, Ipswich, MA) per sample for 30 min. Samples were then flash-frozen in liquid nitrogen and lyophilized overnight using a benchtop freeze-dryer (Labconco, Kansas City, MO). The following day, dry pellet masses were recorded, and pellets were extracted in 10 mL of methanol. Samples were probe-sonicated at 35% intensity with 2 s on / 2 s off pulses for a total of 4 min. Samples were centrifuged at 4,000 × *g* for 6 min, and the supernatants were transferred to new 15 mL conical tubes and dried under nitrogen using a Reacti-Vap evaporator (Thermo Fisher, Waltham, MA). Dried extracts were reconstituted in 250 µL of IPA:ACN:water (8:1:1, v/v/v) and transferred to 1.5 mL microcentrifuge tubes. Extracts were clarified by centrifugation at 10,000 × *g* for 5 min, and the supernatants were transferred to LC/MS vials (Thermo Fisher, Waltham, MA).

### Liquid chromatography high-resolution mass spectrometry (LC-HRMS) quantification by ESI

LC-HRMS analyses were performed on an Agilent 6530 Quadrupole Time-of-Flight (Q-TOF) mass spectrometer coupled to an Agilent 1290 Infinity UHPLC (Agilent, Santa Clara, CA). Separation was carried out on an Agilent InfinityLab Poroshell Aq-C18 column (2.7 µm, 3.0 × 150 mm) maintained at 40 °C, with a flow rate of 0.5 mL $min^{-1}$. Solvent A consisted of water with 0.1% (v/v) formic acid; solvent B consisted of 100% acetonitrile. The gradient began at 100% A and was held for 1 min, with the flow directed to waste for the first 30 s. The flow was then switched to the Q-TOF source, and the gradient ramped linearly from 100% A to 100% B over 10 min, followed by a 30 s hold at 100% B. The column was re-equilibrated to initial conditions for 1 min. The injection volume was 10 µL. Ionization was performed by electrospray ionization (ESI) with a gas temperature of 325 °C, drying gas ($N_2$) flow rate of 4 L $min^{-1}$, and nebulizer pressure of 30 psig. The sheath gas temperature was 325 °C with a sheath gas flow of 11 L $min^{-1}$. The capillary voltage (VCap) was set to 3,500 V, and the nozzle voltage was maintained at 1,000 V. Samples were analyzed in negative polarity in MS1 over a mass range of 100–1,100 *m/z*. Data analysis was performed using MassHunter Qualitative Analysis software (Agilent, Santa Clara, CA).

### Liquid chromatography high-resolution mass spectrometry with data-dependent acquisition (LC-HRMS DDA) by ESI

LC-HRMS analyses were performed on an Agilent 6530 Quadrupole Time-of-Flight (Q-TOF) mass spectrometer coupled to an Agilent 1290 Infinity UHPLC (Agilent, Santa Clara, CA). Separation was carried out on an Agilent InfinityLab Poroshell Aq-C18 column (2.7 µm, 3.0 × 150 mm) maintained at 40 °C, with a flow rate of 0.5 mL $min^{-1}$. Solvent A consisted of water with 0.1% (v/v) formic acid; solvent B consisted of 100% acetonitrile. The gradient began at 100% A and was held for 1 min, with the flow directed to waste for the first 30 s. The flow was then switched to the Q-TOF source, and the gradient ramped linearly from 100% A to 100% B over 10 min, followed by a 30 s hold at 100% B. The column was re-equilibrated to initial conditions for 1 min. The injection volume was 10 µL. Ionization was performed by electrospray ionization (ESI) with data-dependent acquisition (DDA), using a gas temperature of 325 °C, drying gas ($N_2$) flow rate of 4 L $min^{-1}$, and nebulizer pressure of 30 psig. The sheath gas temperature was 325 °C with a sheath gas flow of 11 L $min^{-1}$. The capillary voltage (VCap) was set to 3,500 V, and the nozzle voltage was maintained at 1,000 V. Samples were analyzed in negative polarity over a mass range of 100–1,100 *m/z* in MS1 and 50–1,000 *m/z* in MS2. The collision energy was set to 25 eV, and 5 precursors were selected per cycle. An inclusion list containing 276.9884 *m/z* was used to ensure fragmentation of MEcPP. Data analysis was performed using MassHunter Qualitative Analysis software (Agilent, Santa Clara, CA).

### Liquid chromatography high-resolution mass spectrometry (LC-HRMS) quantification of MK-7 and undecaprenyl species by APCI

LC-HRMS analyses were performed on an Agilent 6530 Quadrupole Time-of-Flight (Q-TOF) mass spectrometer coupled to an Agilent 1290 Infinity UHPLC (Agilent, Santa Clara, CA). Separation was carried out on an Agilent InfinityLab Poroshell EC-C8 column (2.7 µm, 3.0 × 150 mm) maintained at 40 °C, with a flow rate of 0.4 mL $min^{-1}$. Solvent A consisted of 60% acetonitrile and 40% water with 0.1% (v/v) formic acid; solvent B consisted of 90% isopropanol and 10% acetonitrile. The gradient began at 100% A and was held for 1 min, during which the flow was directed to waste. The flow was then switched to the Q-TOF source, and the gradient ramped linearly from 100% A to 100% B over 9 min, followed by a 2 min hold at 100% B. The column was re-equilibrated to initial conditions for 1 min. The injection volume was 5 μL. Ionization was performed by atmospheric pressure chemical ionization (APCI) with a gas temperature of 325 °C, vaporizer temperature of 350 °C, drying gas ($N_2$) flow rate of 3 L $min^{-1}$, and nebulizer pressure of 30 psig. The sheath gas temperature was 325 °C with a sheath gas flow of 11 L $min^{-1}$. The capillary voltage (VCap) was set to 4,000 V, and the corona current was maintained at 6 μA. Samples were analyzed in positive polarity in MS1 over a mass range of 100–1,000 *m/z*. Data analysis was performed using MassHunter Qualitative Analysis software (Agilent, Santa Clara, CA).

### Growth Curve Analyses with *B. subtilis*

We monitored the growth of *B. subtilis* in a 96-well format as follows. In a transparent 96-well round-bottom plate (ThermoFisher, Waltham, MA), 93 μL of LB medium was added to each well, followed by SIE-IPP or SIE-DMAPP from a 100× stock (50 or 100 mM) prepared in dimethyl sulfoxide (DMSO). To confirm that DMSO from these stock solutions did not interfere with the assay, separate wells containing DMSO alone served as controls. Xylose was then added from a 20% stock in ultrapure water to a final concentration of 0.5%. Finally, nucleosides were added from 100× stocks prepared in ultrapure water. Control wells received ultrapure water in volumes equivalent to the xylose wells and, where applicable, nucleoside additions. After all compounds were added, an exponentially growing culture of *B. subtilis* at an $OD_{600}$ of 1 was added to each well at a 1:20 dilution.

The plate was placed into a Cytation 1 cell imaging multimode microplate reader (Agilent, Santa Clara, CA), and $OD_{600}$ measurements were taken every 10 min for 10 h at 30 °C with orbital shaking at 528 cycles per minute (cpm). Growth curves were plotted using a custom R script (14).
